# A comprehensive toolkit for quick and easy visualization of marker proteins, protein-protein interactions and cell morphology in *Marchantia polymorpha*

**DOI:** 10.1101/2020.04.20.050054

**Authors:** Jens Westermann, Eva Koebke, Roswitha Lentz, Martin Hülskamp, Aurélien Boisson-Dernier

## Abstract

Even though stable genomic transformation of sporelings and thalli of *Marchantia polymorpha* is comparatively straightforward and efficient, numerous problems can arise during critical phases of the process such as efficient spore production, poor selection capacity of antibiotics or low transformation efficiency. It is therefore also desirable to establish quick methods not relying on stable transgenics to analyze the localization, interactions and functions of proteins of interest. The introduction of foreign DNA into living cells via biolistic mechanisms has been first reported roughly 30 years ago and has been commonly exploited in established plant model species such as *Arabidopsis thaliana* or *Nicotiana benthamiana*. Here we report the fast and reliable transient biolistic transformation of Marchantia thallus epidermal cells using fluorescent protein fusions. We present a catalogue of fluorescent markers which can be readily used for tagging of a variety of subcellular compartments. Moreover, we report the functionality of the bimolecular fluorescence complementation (BiFC) in *M. polymorpha* with the example of the p-body markers MpDCP1/2. Finally, we provide standard staining procedures for live cell imaging in *M. polymorpha*, applicable to visualize cell boundaries or cellular structures, to complement or support protein localizations and to understand how results gained by transient transformations can be embedded in cell architecture and dynamics. Taken together, we offer a set of easy and quick tools for experiments that aim at understanding subcellular localization, protein-protein interactions and thus functions of proteins of interest in the emerging early diverging land plant model *M. polymorpha*.

## Introduction

In the last decade the liverwort *Marchantia polymorpha* has emerged as a powerful model system to study early land plant evolution due to its early evolutionary divergence in the land plant phylogenetic tree (Shaw et al., 2011; Harrison, 2017; Morris, Puttick et al., 2017). Research deploying *M. polymorpha* has led to a series of insightful studies on the functional evolution of ABA (Lind et al., 2015; Eklund et al., 2018) and JA signaling mechanisms (Monte et al., 2018; Monte et al., 2019; Peñuelas et al., 2019), plant immunity (Carella et al., 2019; Gimenez-Ibanez et al., 2019), reproductive and vegetative development (Flores-Sandoval et al., 2015; Proust et al., 2016; Jones and Dolan, 2017; Rövekamp et al., 2017; Otani et al., 2018; Westermann et al., 2019) and cell division (Buschmann et al., 2015). It offers the advantage of genetic and morphological simplicity in combination with its dominant haploid vegetative life phase, allowing for fast generation of knockout mutants and subsequent phenotypic analyses, irrespectively of time-consuming homozygous mutant generation (Ishizaki et al., 2015 (B)). Concomitantly, a plethora of molecular genetic tools was developed that include stable transformation of developing spores (Ishizaki et al., 2008) and regenerating thallus fragments (Kubota et al., 2013), the suitability for genome editing via homologous recombination (Ishizaki et al., 2013) and CRISPR/Cas9 (Sugano et al., 2014; Sugano et al., 2018), the cultivation in axenic conditions and on soil and controlled crossing (Ishizaki et al., 2015 (B)).

Plant genetics and cell biological approaches generally rely on the efficient visualization of intracellular features, including protein localization and organelle architecture or dynamics. In this regard, the process of transient and stable transformation of plant cells is a powerful and commonly used technique in molecular genetics and cell biology to study protein dynamics, as well as genetic and physical (*i.e.* protein) interaction. It thus aids the elucidation of fundamental biological questions at the (sub)cellular scale. While the performance of stable biolistic transformation of immature thalli and spores has been reported before (Chiyoda et al., 2008, Chiyoda et al., 2014), we describe here the transient biolistic transformation of Marchantia thallus epidermal cells, a technique to study protein localization that has commonly been used in other plant systems for 30 years (Sanford, 1990; Ueki et el., 2009; Rasmussen et al., 1994). Importantly, we provide a comprehensive list of protein marker constructs that allows quick visualization of a variety of subcellular compartments within 24 hours and the possibility for live-imaging. The marker list comprises constructs for visualization of the nucleus, cytoplasm, plasma membrane, actin filaments, endosomes, peroxisomes, the Golgi apparatus and processing bodies (Tab. S1).

Genetic interaction studies often rely on assessment of physical protein interactions to elucidate intracellular signaling mechanisms. Therefore, the bimolecular fluorescence complementation technique (BiFC; Hu et al., 2002; Walter et al., 2004) represents a time-efficient method to test for potentially interacting proteins *in vivo*. Hence, we also provide here evidence for the functionality of BiFC in Marchantia epidermal cells.

In addition to transient expression, dye-based staining procedures represent a fast and reliable method to (co)visualize subcellular compartment architecture and dynamics. Therefore, we here provide a series of staining protocols for different organelles, both for Marchantia thallus epidermal cells and rhizoids and compare functionality regarding standard protocols used for the seed plant model *Arabidopsis thaliana*.

To further complement the here provided comprehensive cell biological marker toolkit, we compiled a list of additional Marchantia resources, methods, tools and databases (Tab. 1) that altogether will be useful for the young and growing community by complementing and supporting genetic / cell biological / biochemical approaches using *M. polymorpha* as a model system.

**Tab. 1:** Important Marchantia resources.

| GENE and GENOME DATABASES |  |  |
| --- | --- | --- |
| Resource/method | Link | Reference |
| Marchantia genome sequence and database | http://marchantia.info/ | Bowman et al., 2017 |
| Marchantia entry on Phytozome (including BLAST and Genome Browser) | https://phytozome.jgi.doe.gov/pz/portal.html#!info?alias=Org_Mpolymorpha | Bowman et al., 2017 |
| Marchantia chloroplast genome studies |  | Ohyama et al., 1988, Umesono et al., 1988, Fukuzawa et al., 1988, Kohchi et al., 1988 |
| MarpoDB : gene-centric database for *Marchantia polymorpha* genetic parts for purposes of genetic engineering and synthetic biology | http://marpodb.io/query | Delmans et al., 2016 |
| PlantTFDB : Plant Transcription Factor Database | http://planttfdb.cbi.pku.edu.cn/index.php?sp=Mpo | Jin et al., 2017<br>Jin et al., 2015<br>Jin et al., 2014 |
| TRANSIENT and STABLE GENETIC MODIFICATION |  |  |
| Homologous recombination-mediated genome editing |  | Ishizaki et al., 2013 |
| Stable Agrobacterium-mediated thallus transformation |  | Kubota et al., 2013 |
| Stable Agrobacterium-mediated sporeling transformation |  | Ishizaki et al., 2008 |
| Design of Gateway-compatible vectors for expression in Marchantia |  | Ishizaki et al., 2015 (A); Mano et al., 2018 |
| CRISPR-Cas-based genome editing |  | Sugano et al., 2014, Sugano et al., 2018 |
| CRISPRdirect target search | https://crispr.dbcls.jp/ | Naito et al., 2015 |
| Stable biolistic thallus transformation |  | Chiyoda et al., 2008 |
| Comprehensive catalogue of fluorescent cell compartment markers |  | This study |
| Protein-protein interaction studies via BiFC |  | This study |

| CELLULAR STAINING TECHNIQUES |  |  |
| --- | --- | --- |
| FM4-64 staining of epidermal cells and rhizoids |  | Kato et al., 2017 (gemmae cups); This study (whole thallus and rhizoids) |
| FM1-43 staining of epidermal cells |  | Minamino et al., 2017 |
| PI staining of thallus epidermal cells and rhizoids |  | Fixed cells: Buschmann et al., 2015; Rövekamp et al., 2016<br>Living cells: Jones and Dolan, 2017; this study |
| DAPI staining of epidermal cells |  | Kondue et al., 2019; this study |
| FDA staining of thallus epidermal cells and rhizoids |  | This study |
| FURTHER RESOURCES |  |  |
| Expressed sequence tags (EST) sequencing |  | Nagai et al, 1999; Nishiyama et al., 2000 |
| RNA sequencing of the gametophyte transcriptome |  | Sharma et al., 2014 |
| 3D imaging using micro-computed tomography and mathematical image-processing method |  | Furuya et al., 2019 |
| Guidelines for Marchantia gene nomenclature |  | Bowman et al., 2015 |

## Materials and Methods

### Plant material and growth conditions

The widely used *Marchantia polymorpha* Tak-1 (MpTak-1) ecotype was cultivated via propagation of vegetative propagules (gemmae) on solid Johnson’s agar supplemented with 0.8 % micro agar under axenic conditions. Gemmae were grown under long day condition (16 h light/8 h darkness cycle) and white light irradiation (60 μmol m^−1^ s^−1^) at 21°C and 60 % humidity. After 2.5 −3 weeks, a few thallus fragments of approximately 0.5 – 1 cm^2^ were transferred onto a small petri dish (6 – 9 cm in diameter) containing fresh solid Johnson’s medium on the day of transformation (Fig. 1A).

**Fig. 1:**
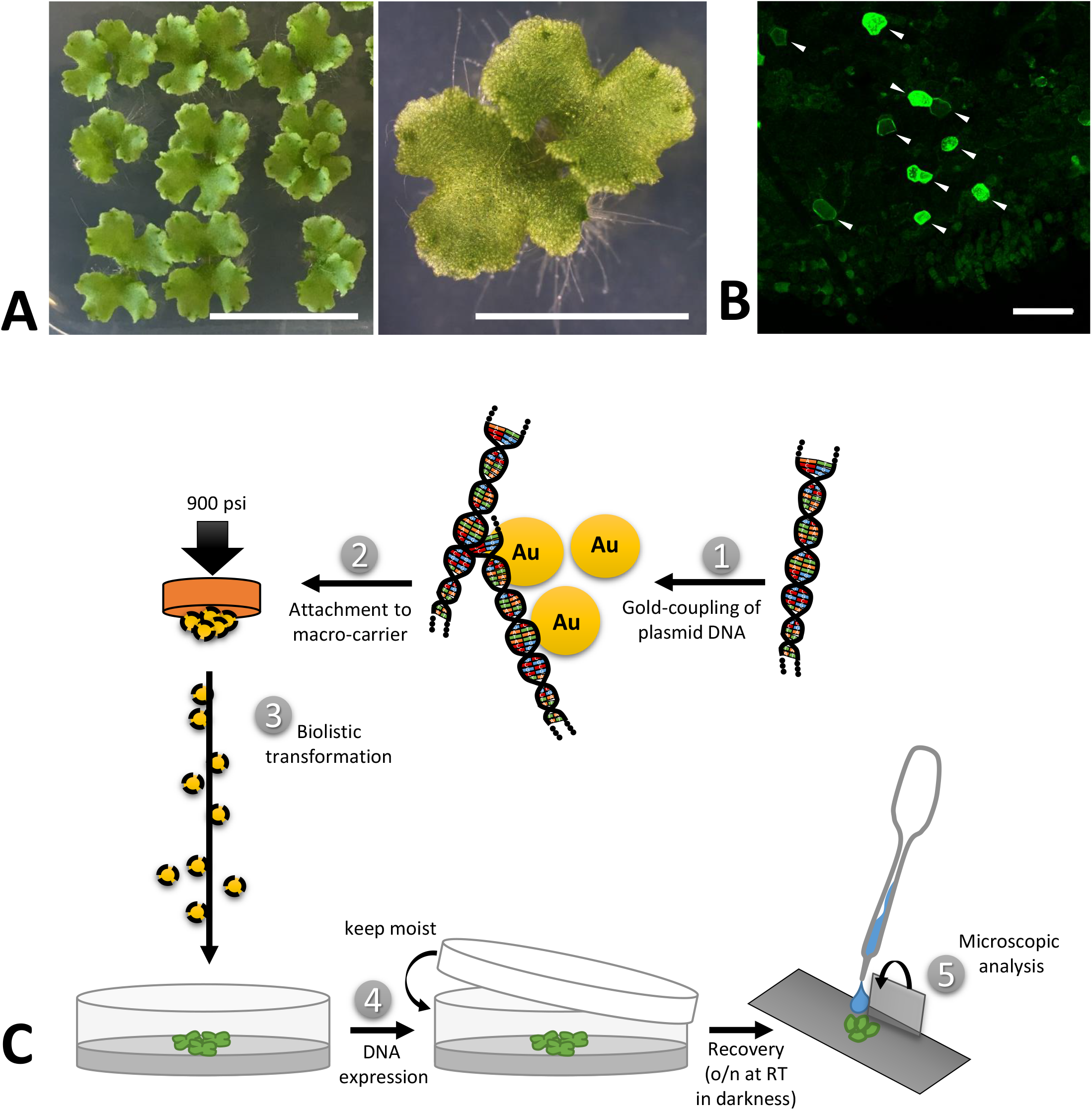
Biolistic transformation of Marchantia thalli. (A) Plant material used for the transformation, showing 2.5 weeks old thalli grown on solid Johnson‘s medium. Scale bars = 2 cm (left) and 5 mm (right). (B) Representative overview of transformation efficiency; arrowheads pointing at transformed cells expressing MpMRI-YFP; Scale bar = 100 μm. (C) Schematic transformation procedure: Vectorial DNA was coupled to gold particles (1), attached to a macro-carrier (2), biolistically transferred into thallus fragments (3), plants were allowed to rest overnight (4) and the pieces expressing the construct of interest were analysed under a microscope (5).

The *Arabidopsis thaliana* Col-0 ecotype, that was used for DAPI staining, was cultivated on soil and grown under long day conditions at 21°C and 120 μmol m^−1^ s^−1^ light intensity.

### Cloning of DNA constructs

All constructs used in this study are summarized in Tab. S1, including their origin, promoter, fluorescent tag and oligonucleotide sequences used for PCR-based amplification of new constructs from Marchantia whole-thallus cDNA. The 35S promoter was used for all expression experiments (except for expression of AtSYP32, AtGot1p and Lifeactin) to guarantee comparability of subsequent analyses. The coding sequences of interest were cloned into Gateway (GW)-compatible entry vectors, pDONR201 and pDONR207 (Invitrogen), and then remobilized to be integrated in the respective GW destination vectors (Tab. S1). The cloning procedure was as described before (Westermann et al., 2019).

### DNA sample preparation for biolistic transformation

For a single shot, 300 ng of vector DNA were mixed with gold, serving as micro-carriers (30 mg/ml, 1 μm), CaCl_2_ (2.5 M), spermidine (0.1 M) and _dd_H_2_O under thorough shaking. Subsequently, micro-carriers were washed with 70% EtOH and 100% EtOH. The DNA-coated gold particles were suspended in 100% EtOH and placed onto macro-carriers. The EtOH was allowed to vaporize and the prepared macro-carriers were then used for biolistic transformation.

### Biolistic transformation procedure and efficiency of (co-)transformation

Marchantia thallus fragments were placed into a PDS-1000 / He Biolistic^®^ Particle Delivery System (Bio-Rad). A vacuum of 25 in Hg vac was applied and the DNA-coated gold particles were shot at 900 psi from a distance of 10 cm. Finally, the bombarded plant material was allowed to recover for 24 h in darkness while remaining in its humid environment, *i.e.* on the media in the closed petri dish (Fig. 1C). Biolistic transformation generally yielded n > 50 transformed cells per sample shot. A representative example for transformation efficiency is shown in Fig. 1B. Moderate to strong expression levels in each individual cell could be observed irrespectively of the protein construct or promoter used (Fig. 1B; Tab. S1). The use of strong promoters such as pro35S, proAtUBQ10 or proMpEF1α can sometimes lead to overexpression artefacts that may impede drawing secured conclusions. However, yielding a wide range of expression level in the same experimental round and plant sample allows for identification of biologically meaningful protein localization patterns and to distinguish them from unwanted artefacts, such as protein over-accumulations. In order to reliably assess the potential of transformed constructs as single cell fluorescent markers, we co-bombarded all described vectors with either the nuclear marker AtKRP1 or the plasma membrane markers AtNPSN12 or MpSYP13a fused to fluorescent tags and subsequently created a collection of functional and useful Arabidopsis- and Marchantia-derived fluorescent protein fusions (Tab. S1). In order to determine the efficiency of co-transformation, we counted cells expressing both markers in relation to the total number of transformed cells in 9 independent co-transformations of protein fusions used in this study. Successful biolistic co-transformation was on average 74% efficient (see Tab. S3).

### Staining procedures

For FDA staining, young (2 – 5 days-old) gemmae were placed onto depression slides and covered with an FDA solution (5mg/L FDA in _dd_H_2_O, diluted from a stock solution of 5 mg/ml FDA in acetone) for 5-10 min. Afterwards, samples were rinsed in _dd_H_2_O.

For PI staining, young gemmae were placed onto depression slides and directly covered with a PI solution for 10 minutes (10 mg/L in _dd_H_2_O). Subsequently, samples were rinsed with _dd_H_2_O.

For FM4-64 staining, young gemmae were mounted onto depression slides in 2 μM FM4-64 diluted in liquid Johnson’s growth medium (Ishizaki et al., 2008) and allowed to incubate for 10 minutes prior to imaging. For FM4-64 and FDA co-staining, gemmae were first stained in a FDA solution and then mounted in a FM4-64 solution, both as described above.

For DAPI staining, several methods were used. Experiments were done using 0, 4, and 7 days old gemmae. The DAPI staining solutions were composed of 10 – 100 mg/L DAPI in either 1xPBS-T (phosphate buffered saline + 0.1% Tween-20) and 5% DMSO or _dd_H_2_O with 0.1 or 1% Tween-20 and 5% DMSO. Different staining incubation times of 10, 30 or 60 minutes were tested. The staining was tested with and without preceding or subsequent shaking of the samples in 70% EtOH at 80°C. To enhance permeability of membranes, 10 or 50 mg/L digitonin was added to the aforementioned staining solutions. As all attempts for staining living cells failed, the following fixation methods were tested. Gemmae were fixed in a 3:1 EtOH:acetic acid mixture on ice for 1 h, washed 3 times in 100% EtOH and stained in aforementioned DAPI solutions for 1 h. In another attempt, gemmae were fixed in 3% glutaraldehyde in 1x PBS-T (phosphate buffered saline + 0.1% Tween-20) overnight, subsequently washed in 1x PBS-T, and incubated in aforementioned DAPI solutions in darkness overnight. Furthermore, a modified version of a DAPI staining protocol published for gametophore leaves and protonemata of *Physcomitrella patens* (Sato et al., 2017) was used. Gemmae were placed in 3.7% formaldehyde in 1 x PBS for 30 min. Subsequently gemmae were immersed in 100% MeOH on ice for 10 min. Afterwards, gemmae were soaked in 1% Triton X-100 and then stained with the aforementioned DAPI solutions for 30 min. Unfortunately, none of the experimental procedures described here led to a reliable staining of nuclei by DAPI in viable or dead epidermal cells of *M. polymorpha* gemmae.

### Confocal laser scanning microscopy

The transformed or stained plant material was transferred onto a depression slide supplemented with 300 μL _dd_H_2_O and covered with a 18×18 mm cover slip. Microscopic analysis was performed using a Leica SP8 CLSM with an argon gas laser intensity set at 20 %. Fluorophore excitation and fluorescence caption were performed at the wavelength spectra shown in Tab. S2. Images were taken using a digital gain of 100 % at a resolution of 1024 x 1024 pixels, a pinhole size of 1 AU, and a scan speed of 400-700 Hz using bidirectional confocal scanning and HyD hybrid detectors. For the caption of multiple fluorophore types sequential or, if suitable, simultaneous scanning was performed.

### Data processing and analysis

Analysis of all microscopic captions was performed using ImageJ/FIJI (Schindelin et al., 2012), software version 1.51n. Data manipulation included maximum projections from Z-stacks (≤ 20 frames, 1 μm slice intervals) for some of the markers (as individually mentioned in the figure captions), as well as generation of composite images from separate individual channels.

## Results

### A) Fluorescent protein markers to illuminate cellular compartments in Marchantia

To assess the potential capability of transiently transforming single Marchantia thallus epidermal cells, we first chose a set of proteins whose subcellular localization has been well studied in established model systems such as Arabidopsis or tobacco and thus could qualify as reliable subcellular markers in Marchantia as well.

#### Nucleus

We first picked the *Arabidopsis thaliana* INHIBITOR OF CYCLIN-DEPENDENT KINASE 1 (AtICK1)/ KIP-RELATED PROTEIN 1 (AtKRP1), which localizes to the nucleus and functions in cell growth, differentiation and cell cycle progression (Wang et al., 1998; De Veylder et al., 2001; Schnittger et al., 2003; Weinl et al., 2005; Jakoby et al., 2006). Upon biolistic transformation of Marchantia thalli, we observed AtKRP1-CFP protein localization to the nucleus of epidermal cells (Fig. 2A). We therefore co-transformed AtKRP1 as a nuclear marker and indicator of successful cell transformation in subsequent experiments.

**Fig. 2:**
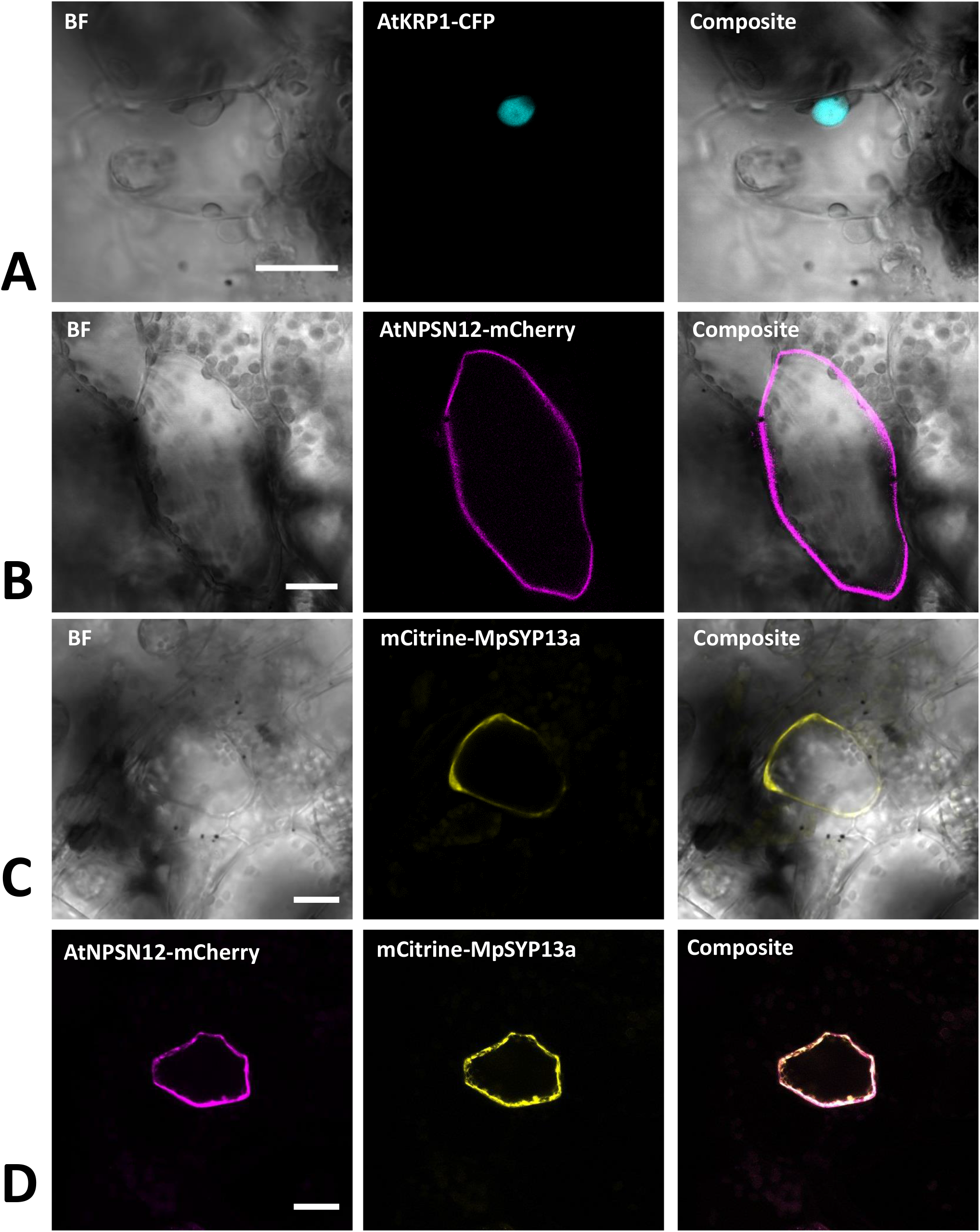
Confirmation of known nuclear and plasma membrane markers. (A) The Arabidopsis nuclear marker AtKRP1 localizes to the nucleus of *M. polymorpha* epidermal cells. (B) The Arabidopsis plasma membrane marker AtNPSN12 localizes to the plasma membrane in Marchantia epidermal cells. (C, D) The Marchantia plasma membrane marker MpSYP13a co-localizes with AtNPSN12. All scale bars = 20 μm. BF = bright field.

##### Plasma membrane

As a second potential marker, we chose the *Arabidopsis thaliana* NOVEL PLANT SNARE 12 (AtNPSN12), which represents a non-polar plasma membrane-localized protein commonly used as plasma membrane marker (Alassimone et al., 2012; Kirchhelle et al., 2016). Biolistically transformed Marchantia thallus epidermal cells showed AtNPSN12-mCherry fluorescence at the cell periphery consistent with plasma membrane localization (Fig. 2B). To confirm this localization, we co-transformed AtNPSN12-mCherry with the known Marchantia plasma membrane marker mCitrine-MpSYP13a (Kanazawa et al., 2016; Fig. 2C). As single and co-bombardments with AtNPSN12 showed (co)localization to the plasma membrane, we conclude that AtNPSN12-mCherry and mCitrine-MpSYP13a are both suitable plasma membrane markers for Marchantia epidermal cells (Fig. 2D).

Receptor-like kinases of the Malectin-like receptor kinase subfamily have been the subject of intensive research in the past years given their multitude of functions in plant development and immunity signaling (Franck et al., 2018 A). The Malectin-like receptors (MLRs) ANXUR1 and 2 (AtANX1/2) control cell wall integrity during pollen tube growth (Boisson-Dernier et al., 2009; Miyazaki et al., 2009) and negatively regulate plant immune responses in Arabidopsis (Mang et al., 2017) at the plasma membrane. During pollen tube growth control, AtANX1/2 act genetically upstream of the cytosolic and plasma membrane-attached receptor-like cytoplasmic kinase of the PTI1-like family, AtMRI, while the AtANX1 homolog AtFERONIA (AtFER) acts upstream of AtMRI during root hair growth control (Boisson-Dernier et al., 2015). We showed recently that tip-growth control in Marchantia rhizoids relies on an evolutionarily conserved signaling module comprised of the unique Marchantia MLR MpFER and its downstream component and unique Marchantia PTI1-like MpMRI, both of which show plasma membrane-localization comparable to their respective Arabidopsis homologs (Westermann et al., 2019). We transiently co-expressed the fluorescent protein fusions AtMRI-YFP, MpFER-YFP and MpMRI-YFP with AtNPSN12-mCherry. While MpFER-YFP showed signal exclusive to the plasma membrane, AtMRI and MpMRI displayed plasma membrane localization with traces in the cytoplasm as reported before (Fig. 3; Boisson-Dernier et al., 2015; Westermann et al., 2019).

**Fig. 3:**
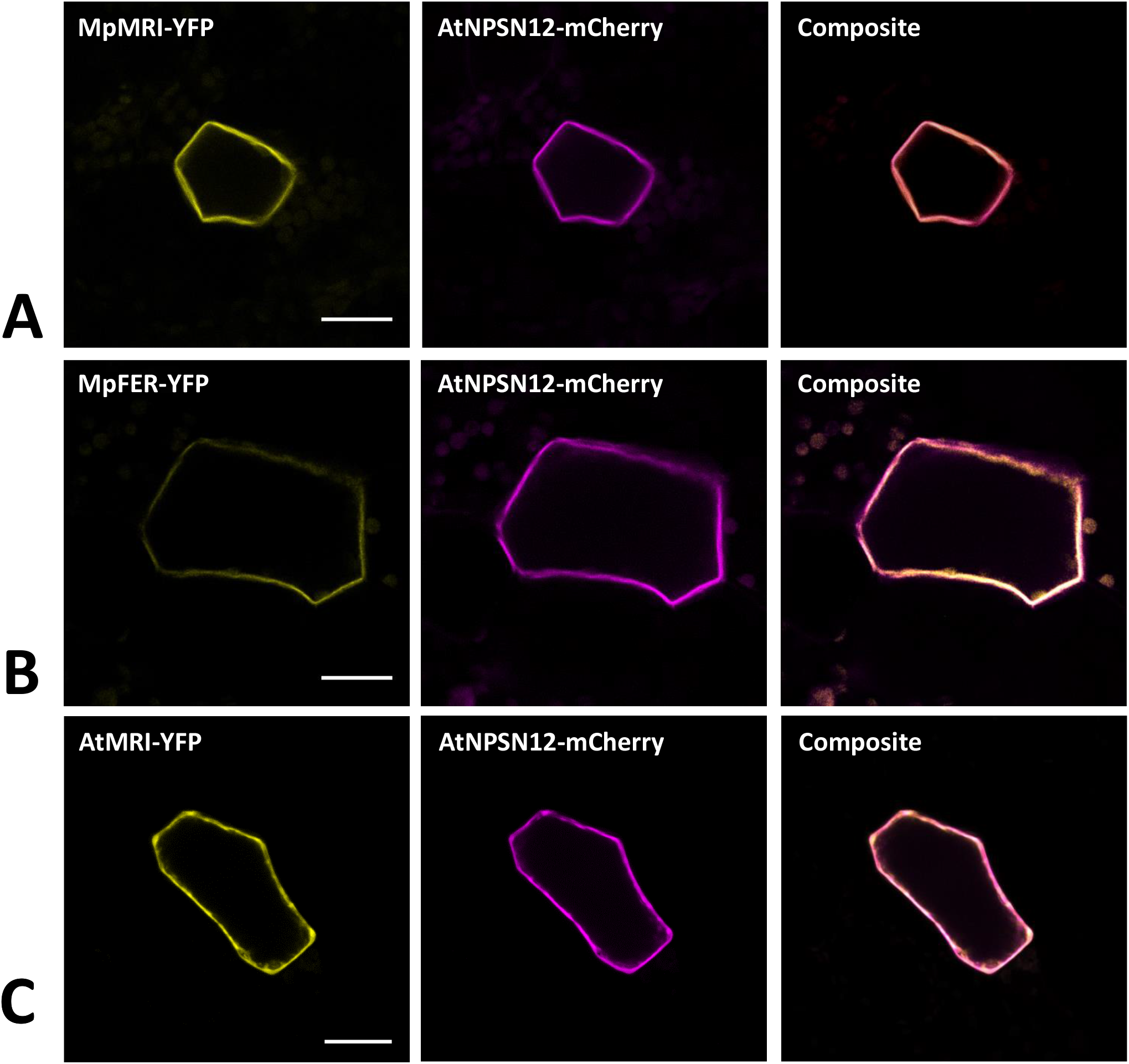
Plasma membrane markers for Marchantia research. MpMRI (A), MpFER (B) and AtMRI (C) all localized to the plasma membrane of *M. polymorpha* thallus epidermal cells. All three constructs co-localized with the plasma membrane marker AtNPSN12. All scale bars = 20 μm.

Noteworthily, we also wanted to test expressing the plasma membrane localized Arabidopsis MLRs in Marchantia and thus co-transformed AtANX1-RFP with mCitrine-MpSYP13a and AtFER-Citrine with AtNPSN12-mCherry. Intriguingly, while many cells expressed the plasma membrane markers mCitrine-MpSYP13a and AtNPSN12-mCherry, a great majority of them did not show expression of either AtANX1 or AtFER (Fig. S1A, B). This suggests that, unlike MpFER, the Arabidopsis MLRs fused to single fluorescent tag cannot be expressed in Marchantia epidermal cells. Thus, we next tried to express AtFER with a triple Citrine tag instead of a single one. It resulted in many Citrine-expressing cells but mostly in the cytoplasm, with no hints of plasma membrane localization (Fig. S1C). These results indicate that fusion of long protein tags may prevent transmembrane receptor kinases such as MLRs to be correctly integrated into cellular membranes. To check if this was specifically due to Arabidopsis proteins or to certain protein families, we co-expressed MpFER-3xCitrine with MpFER-TdTomato and MpMRI-3xCitrine with MpMRI-RFP (Fig. S1D, E). Interestingly, the 3xCitrine tag didn’t perturbate the cytosolic and plasma membrane localization of MpMRI, as MpMRI-3xCitrine co-localized with MpMRI-RFP at the cell periphery. However, while MpFER-TdTomato exhibited PM localization, MpFER-3xCitrine-derived signal was clearly present in the cytoplasm. Therefore, for some plasma membrane-localized protein families, fusion with a triple tag can lead to localization artefacts, and the use of single tag is thus recommended by default. Why MpFER but neither AtFER nor AtANX1 can be expressed in Marchantia thallus epidermis remains puzzling.

##### Cytoplasm

The *A. thaliana* type-one protein phosphatases (TOPP) AtATUNIS1/2 have recently been reported as negative regulators of cell wall integrity maintenance during Arabidopsis tip-growth (Franck et al., 2018 B). The nucleocytoplasmic localization of AtAUN1-YFP and AtAUN2-YFP was demonstrated in Arabidopsis pollen tubes and leaf epidermal cells (Franck et al., 2018 B). In Marchantia epidermal cells, expression of AtAUN1/2-YFP led to a comparable nucleocytoplasmic localization, as opposed to the co-expressed plasma membrane localized AtNPSN12-mCherry fusion (Fig. 4), therefore qualifying these phosphatases as reliable Marchantia nucleocytoplasmic markers.

**Fig. 4:**
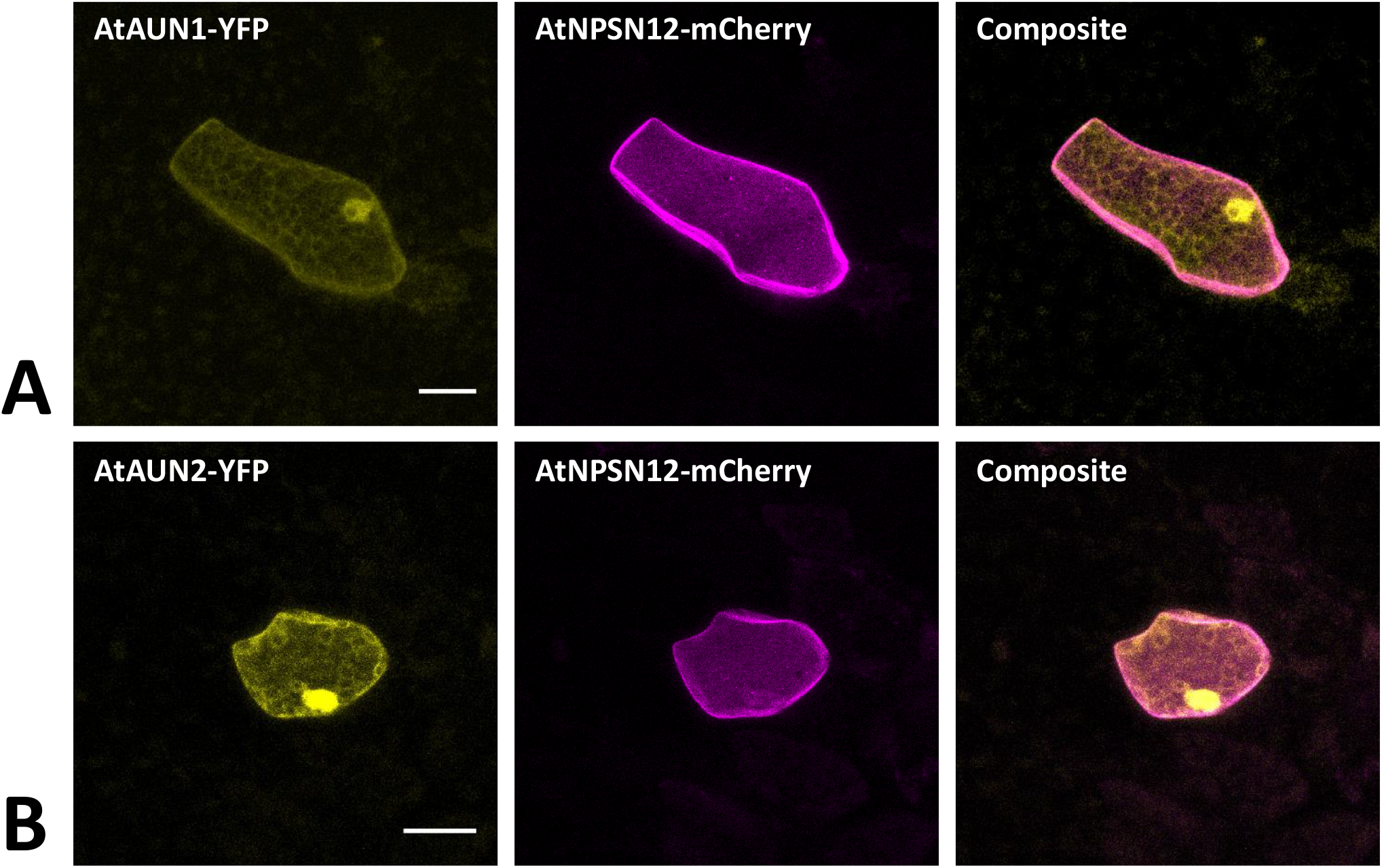
Nucleocytoplasm markers for Marchantia research. Both, AtAUN1 and AtAUN2 localized to the cytoplasm and nucleus of *M. polymorpha* thallus epidermal cells, consistent with observations in *A. thaliana* (Franck et al., 2018). The constructs were co-bombarded with plasma membrane marker AtNPSN12. All scale bars = 20 μm. Pictures show maximum projections of z-stack captions, hence the appearance of the ‘cytoplasmic noise’ signal for AtNPSN12-mcherry (see Materials and methods section for details).

##### Endosomes

As for endosomal compartments, we chose two Ras-related in brain (RAB) GTPases, the canonical MpRAB5 and the plant-unique MpARA6, that were recently described in *M. polymorpha*. Both proteins were successfully expressed in stably transformed lines and co-localized to endosomal punctate structures stained by FM1-43 (Minamino et al., 2017). Upon biolistic co-transformation of the protein fusions mCherry-MpRAB5 and MpARA6-YFP with the nuclear marker AtKRP1-CFP (Fig. 5A and B), we found a comparable localization in punctate structures for both markers. Moreover, both GTPases strongly co-localized with each other (Fig. 5C) showing that MpRAB5 and MpARA6 are suitable endosomal markers also for transient transformation studies.

**Fig. 5:**
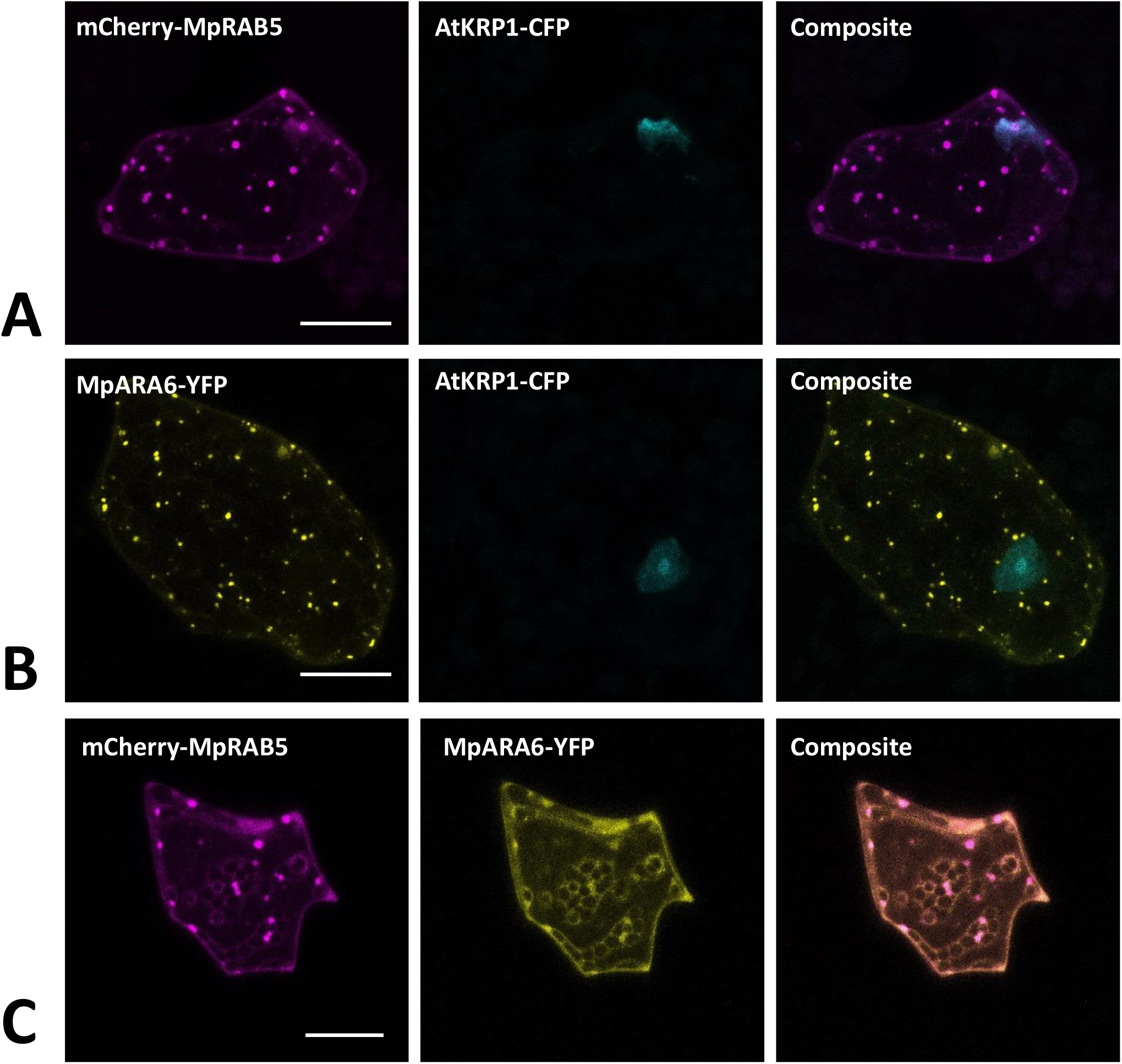
Endosomal markers for Marchantia research. Both, MpRAB5 (A) and MpARA6 (B) localized to punctuate intracellular structures of *M. polymorpha* thallus epidermal cells, likely representing endosomes. The constructs were co-bombarded with nuclear marker AtKRP1. The endosomal markers MpRAB5 and MpARA6 also show clear co-localization (C). All scale bars = 20 μm. Pictures show maximum projections of z-stack captions (see Materials and methods section for details).

##### Peroxisomes

The carboxyl-terminal amino acid sequence serine-lysine-leucine (SKL) is well known as the consensus peroxisomal targeting sequence 1 (PTS1) and is sufficient to induce protein targeting and import to peroxisomes. SKL was first shown to be able to signal protein import into peroxisomes of mammalian cells (Gould et al., 1989) but later was also found to be functional in yeast and plants (Keller et al., 1991). In Arabidopsis, SKL motif fused to fluorescent tags is frequently used as a peroxisomal marker (Mathur et al., 2002, Rodríguez-Serrano et al., 2009, Kim et al., 2013). In *M. polymorpha*, SKL targeting was utilized for evaluation of CRISPR-Cas9 modules (Konno et al., 2018). In transiently transformed *M. polymorpha* cells, we also found a clear and distinct localization of mCherry-SKL in punctate structures, likely representing peroxisomes (Fig. 6A).

**Fig. 6:**
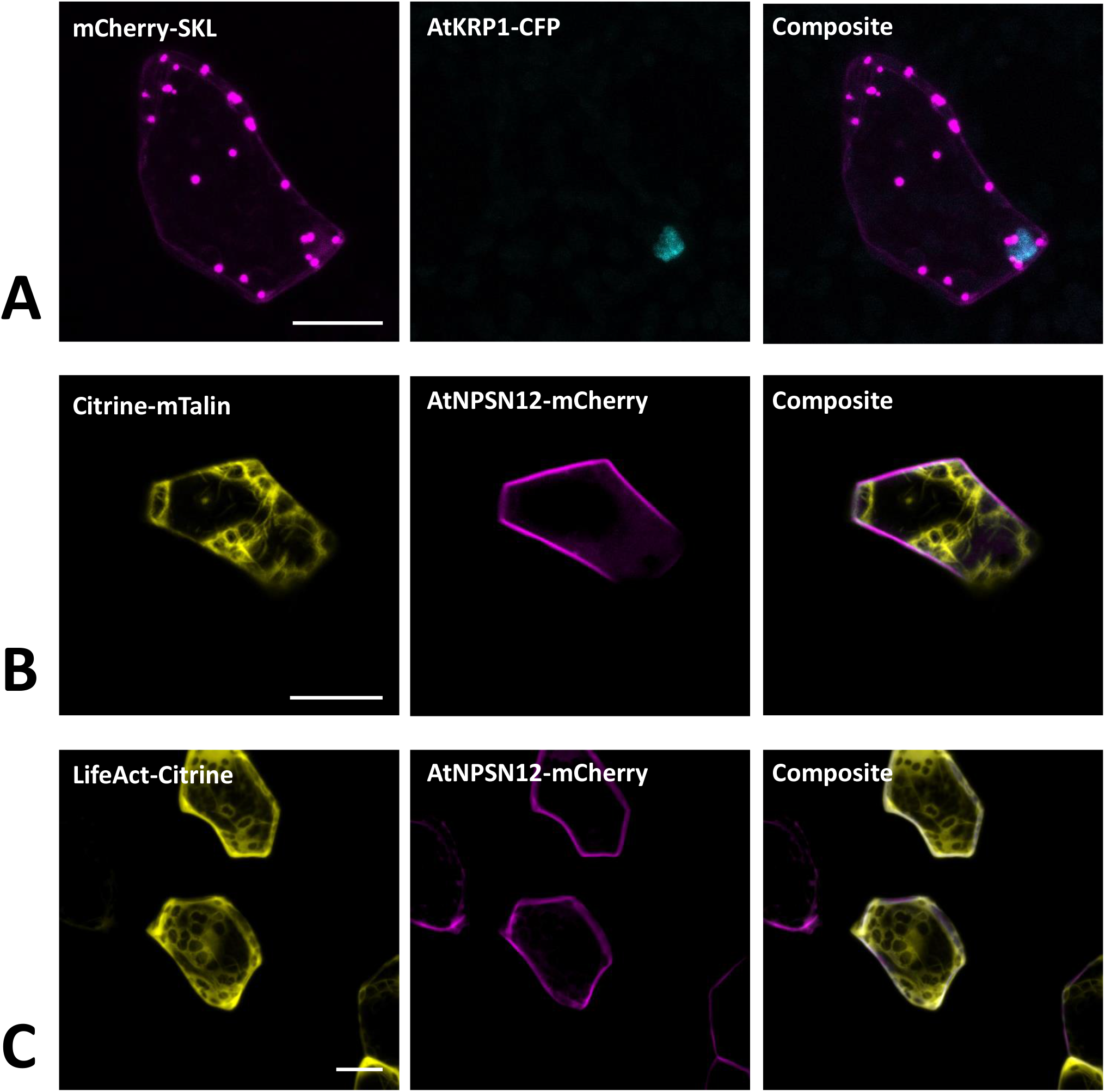
Peroxisomal and actin filaments markers for Marchantia research. (A) The SKL-target sequence tagged to mCherry localized to single intracellular foci of *M. polymorpha* thallus epidermal cells, likely representing peroxisomes. mCherry-SKL was co-bombarded with nuclear marker AtKRP1. The actin filament markers (B) Citrine-mTalin and (C) LifeAct-Citrine were co-bombarded with plasma membrane marker AtNPSN12-Mcherry. All scale bars = 20 μm. Pictures show maximum projections of z-stack captions (see Materials and methods section for details).

##### Actin filaments

Both the Lifeact peptide - a short peptide of 17 amino acids – and the C-terminal 197 amino acids of mouse talin are known to bind to filamentous actin (Kost et al., 1998; Riedl et al., 2008). Therefore, to visualize the actin filaments in Marchantia epidermal cells, we used the Citrine-mTalin and LifeAct-Citrine reporters described previously (Kimura and Kodama, 2016). As in stably transformed Marchantia lines (Kimura and Kodama, 2016), both markers successfully revealed the actin filament networks around chloroplasts in epidermal cells (Fig. 6B, C).

##### Golgi apparatus

As potential markers for the Golgi apparatus, we selected the Arabidopsis proteins SYNTAXIN OF PLANTS 3 (AtSYP3) and the GOLGI TRANSPORT 1 p homolog (AtGot1p). Both proteins have been shown to localize to the Golgi apparatus (Uemura et al., 2004; Conchon et al., 1999) and are reliable Golgi markers for Arabidopsis, as being part of the Wave line multicolor marker set for membrane compartments (WAVE22 and WAVE18, respectively; Geldner et al., 2009). Upon transient biolistic expression of YFP-AtSYP3 and YFP-AtGot1p in *M. polymorpha*, a distinct and comparable localization pattern of both proteins, likely representing the Golgi apparatus, was visible (Fig. 7A and B). Furthermore, upon co-expression of CFP-AtSYP3 and YFP-AtGOT1p, we also found perfect co-localization (Fig. 7C) confirming that both markers are reliable to illuminate the Golgi in Marchantia.

**Fig. 7:**
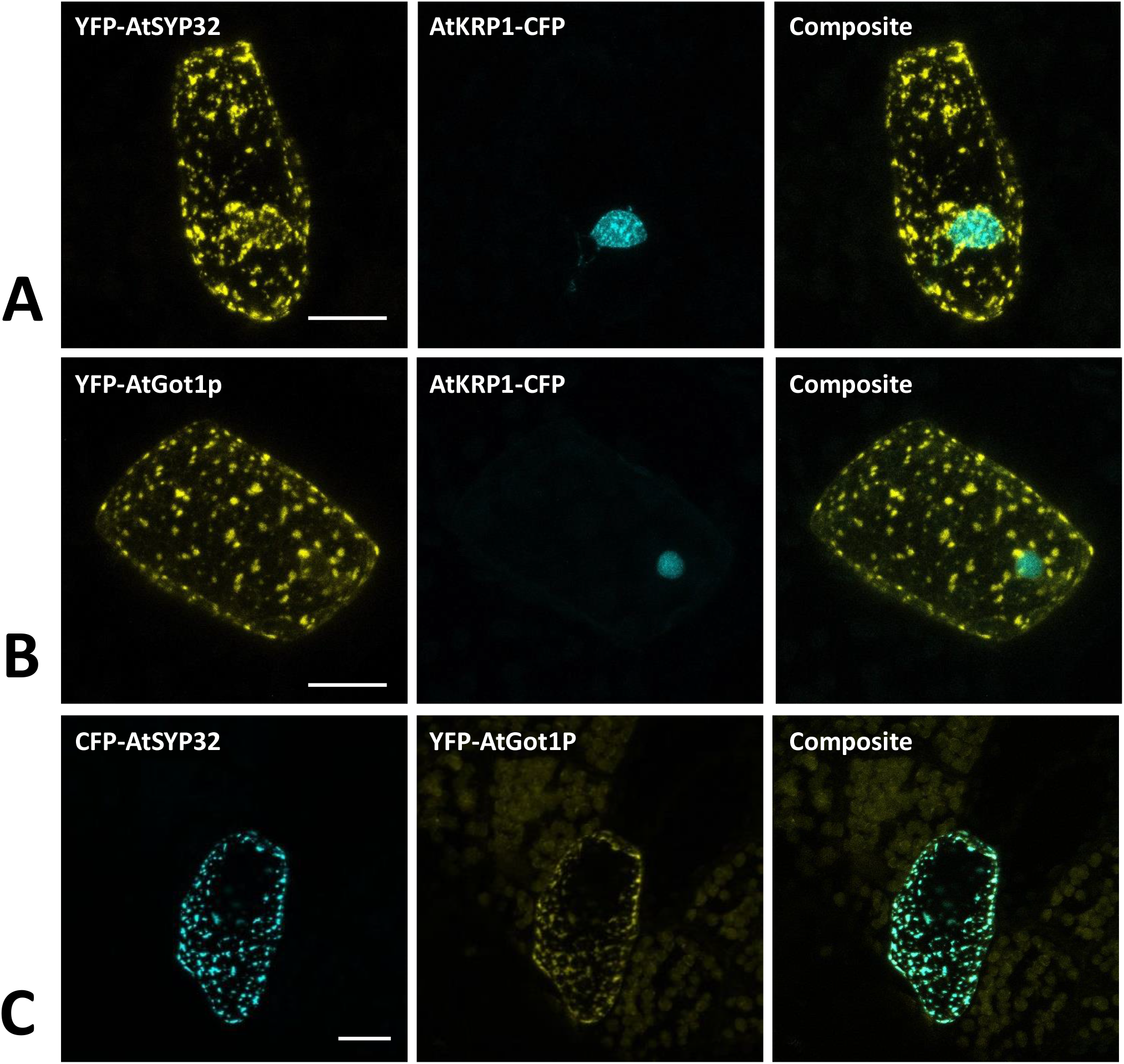
Golgi markers for Marchantia research. The Arabidopsis golgi markers AtGot1p (A) and AtSYP32 (B) localize to the golgi apparatus of *M. polymorpha* epidermal cells. The constructs were co-bombarded with nuclear marker *At*KRP1. (C) The golgi markers AtGot1P and AtSYP32 show clear co-localization. All scale bars = 20 μm. BF = bright field.

##### mRNA processing bodies

mRNA processing bodies (p-bodies), have been found to play a crucial role in mRNA processing comprising deadenylation, decapping, degradation, mRNA storage and mRNA quality control (thoroughly reviewed for *A. thaliana* in Maldonado-Bonilla, 2014). As p-bodies markers, we chose the *Arabidopsis thaliana* DECAPPING PROTEIN 1 (AtDCP1) and AtDCP2, whose function has been well studied in the past years (Xu et al., 2006; Xu and Chua, 2009; Steffens et al., 2015; Bhasin et al., 2017). Upon transformation of the protein fusion AtDCP1-mCherry we found a comparable expression in dot-like structures, likely representing p-bodies (Fig. 8A). In contrast, transformation of mCherry-AtDCP2 revealed a diffused expression throughout the cytoplasm and in the nucleus (Fig. 8B), as reported in Arabidopsis in the absence of stress (Motomura et al., 2014). Therefore, we assume that AtDCP2 is also generally localized in the cytoplasm and nucleus in *M. polymorpha* and is only recruited to p-bodies upon stress conditions (Motomura et al., 2014).

**Fig. 8:**
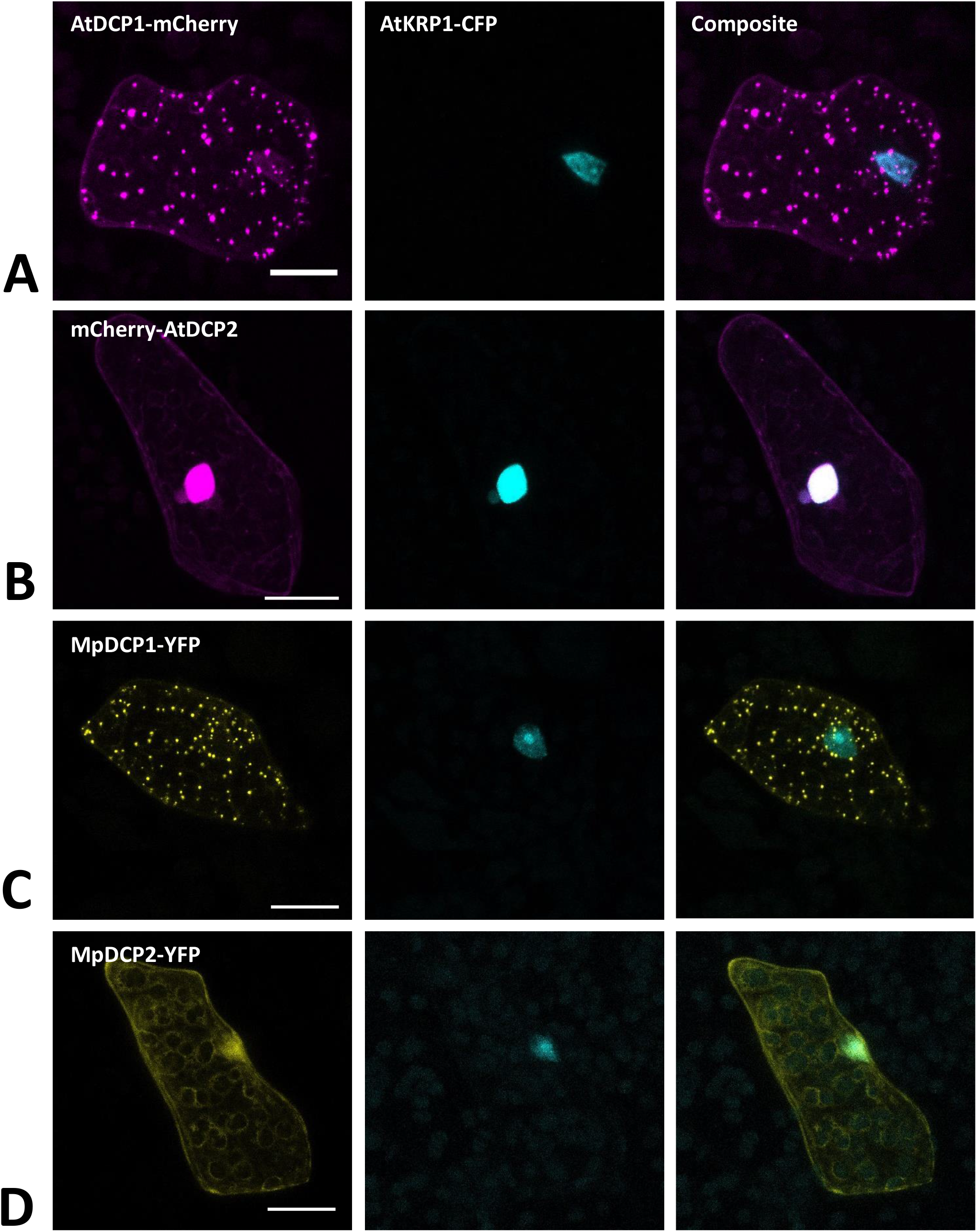
Marchantia p-bodies markers. Both, AtDCP1 (A) and MpDCP1 (C) localized to intracellular dot-like structures, that likely represent p-bodies. In contrast, AtDCP2 (B) and MpDCP2 (D) localized to the cytoplasm, consistent with former observations (Motomura et al., 2014). Additionally, both, AtDCP2 and MpDCP2 showed a nuclear localization, co-localizing with the nuclear signal of AtKRP1. The constructs were co-bombarded with nuclear marker AtKRP1. Scale bar = 20 μm. Pictures show maximum projections of z-stack captions (see Materials and methods section for details).

The similar localization of AtDCP1/2 in Arabidopsis (Iwasaki et al., 2007; Motomura et al., 2014) and Marchantia suggests that the function of DCPs in mRNA processing has been evolutionarily conserved. To assess whether the two Marchantia DCP-homologs MpDCP1/2 localize similarly as their Arabidopsis counterparts, we transformed different combinations of fluorescent fusions (MpDCP1-mCherry, MpDCP1-YFP, MpDCP2-mCherry, MpDCP2-YFP). As anticipated, MpDCP1 displayed a dot-like localization pattern similar to AtDCP1, while MpDCP2 exhibited an AtDCP2-like nucleocytoplasmic localization (Fig. 8C and D).

### B) Bimolecular fluorescence complementation

Based on former reports of AtDCP1 to regulate mRNA decay and to recruit further functionally relevant proteins, such as AtDCP2, to p-bodies (Iwasaki et al., 2007; Motomura et al., 2014), as well as our own observations (see above), we selected MpDCP1/2 as promising candidates to assess the feasibility of studying protein-protein interactions in *M. polymorpha* via bimolecular fluorescence complementation (BiFC). The BiFC technique relies on the co-expression of two proteins fused to the N- or C-terminal part of a fluorescent reporter (*e.g.*-YFP_N_ and -YFP_C_, respectively. Upon physical interaction of the two tagged proteins of interest, the N- and C-terminal parts of the reporter can reconstitute a functional fluorescent protein. Capture of the respective fluorescent signal thus is used as an indicator for protein-protein interaction. For BiFC to be meaningful, the co-transformation of both reporter halves must lead to regular and frequent co-expression, which is the case for Marchantia thallus transient biolistic transformation as it reaches, in our hands, 74% on average (see Material and Methods section and Tab. S3). The physical interaction of AtDCP1/2 was foremost reported in *in vitro* pull-down assays (Xu et al., 2006) and later independently confirmed by BiFC in tobacco mesophyll protoplasts (Weber et al., 2008).

Interestingly, upon co-expression of YFP_N_-MpDCP2 and YFP_C_-MpDCP1, together with the nuclear marker AtKRP1-CFP, we could observe a clear and specific YFP signal in dot-like structures, suggesting that MpDCP1/2 are capable of interacting physically in p-bodies of Marchantia epidermal cells (Fig. 9A). To exclude the possibility of false positive signals (Kodama et al., 2012) in our experimental setup we also transformed YFP_N_-MpDCP2 and YFP_C_-MpDCP1 with YFP_C_–MpLYST interacting protein 5 (MpLIP5) and AtMYC related protein1 (AtMYC1)-YFP_N_ tags, respectively. Expression of both vector combinations led to the absence of a YFP signals in cells expressing AtKRP1-CFP (Fig. 9B and C), indicating that the observed interaction between MpDCP1 and MpDCP2 is specific. The integrity of YFP_C_–MpLIP5 was confirmed by co-expression with the Marchantia homolog of a known interactor of LIP5 in Arabidopsis - MpSuppressor of K^+^ Transport Growth Defect1 (MpSKD1) (Haas et al., 2007), N-terminally fused to YFP_N,_ showing a clear YFP signal in punctate structures consistent with localization to p-bodies (Fig. S2A). The integrity of AtMYC1-YFP_N_ was shown by a BiFC interaction in the nucleus with its known interaction partner AtTRANSPARENT TESTA GLABRA1 (TTG1) (Zimmermann et al., 2014, Zhao et al., 2012; control used in Steffens et al., 2017), C-terminally fused to YFP_C_ (Fig. S2B). In conclusion, our results show that BiFC is functional in Marchantia and can be used to quickly assess protein-protein interactions *in vivo*.

**Fig. 9:**
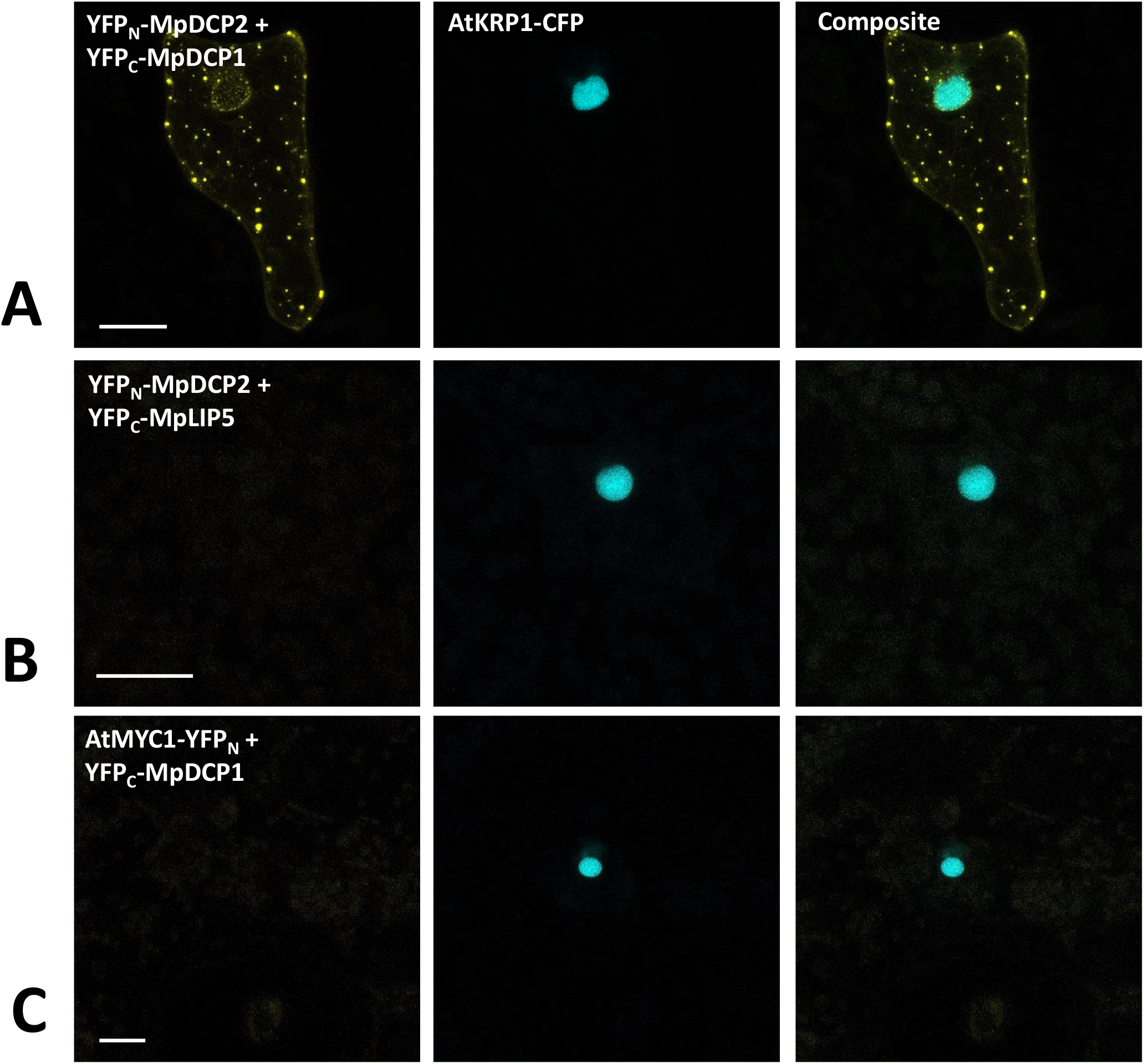
Bimolecular fluorescent complementation assays showing interaction between MpDCP1 and MpDCP2. (A): Co-transformation of split-YFP fusion constructs of MpDCP1 and MpDCP2 result in a fluorescence signal in dot-like foci, indicating protein-protein interaction in p-bodies. (B) Co-bombardment of split-fusions with MpDCP2 and the unrelated MpLIP5 protein were used as a negative control. (C) We also co-bombarded split-versions of MpDCP1 and unrelated AtMYC1, which also led to the absence of any fluorescence signal. The constructs were co-bombarded with nuclear marker AtKRP1. Scale bar = 20 μm. Pictures show maximum projections of z-stack captions (see Materials and methods section for details). See also Fig. S2 for other controls.

### C) Staining intracellular structures in *M. polymorpha*

For the visualization of a cell and the investigation of cellular architecture and dynamics, it is crucial to have several quick and reliable staining methods for live cell imaging at hand. Therefore, we tested some standard staining procedures to label intracellular compartments and cellular structures (including the plasma membrane, cytoplasm, cell wall and nucleus) in *M. polymorpha* gemmae that have been established for other plants but lacking ready-to-use protocols for Marchantia.

#### Fluorescein diacetate (FDA) for cytoplasm staining of living cells

FDA is a cell-permeable, per se non-fluorescent esterase substrate. As soon as it passes the plasma membrane, it is hydrolyzed by esterases in the cytoplasm of viable cells (Rotman and Papermaster, 1966). Thereby, FDA is converted to a negatively charged, green-fluorescent fluorescein unable to either cross back the plasma membrane or pass the tonoplast and thus it accumulates in the cytoplasm. Owing to these properties, FDA is suitable for cell viability assays and can be used as a negative stain for vacuoles. In Arabidopsis, FDA staining has been reliably used for testing root hair and guard cell viability (Schapire at al., 2008; Hao et al., 2012),to visualize vacuoles in root hairs (Saedler et al., 2009) and trichomes (Mathur et al., 2003), and to study pathogen response (Jones et al., 2016).

Here, we successfully utilized FDA to stain the cytoplasm of rhizoids and epidermal cells in young gemmae (Fig. 10). FDA showed a strong, green fluorescence already after a short incubation time of 10 minutes, demonstrating the viability of rhizoids and epidermal cells. We here present FDA as a tool to be readily used for visualization of the cytoplasm in *M. polymorpha*. As it is not able to pass the tonoplast, it can also be used to detect vacuolar architecture, especially in rhizoids, where vacuolar volume was clearly visible after staining with FDA (Fig. 10D).

**Fig. 10:**
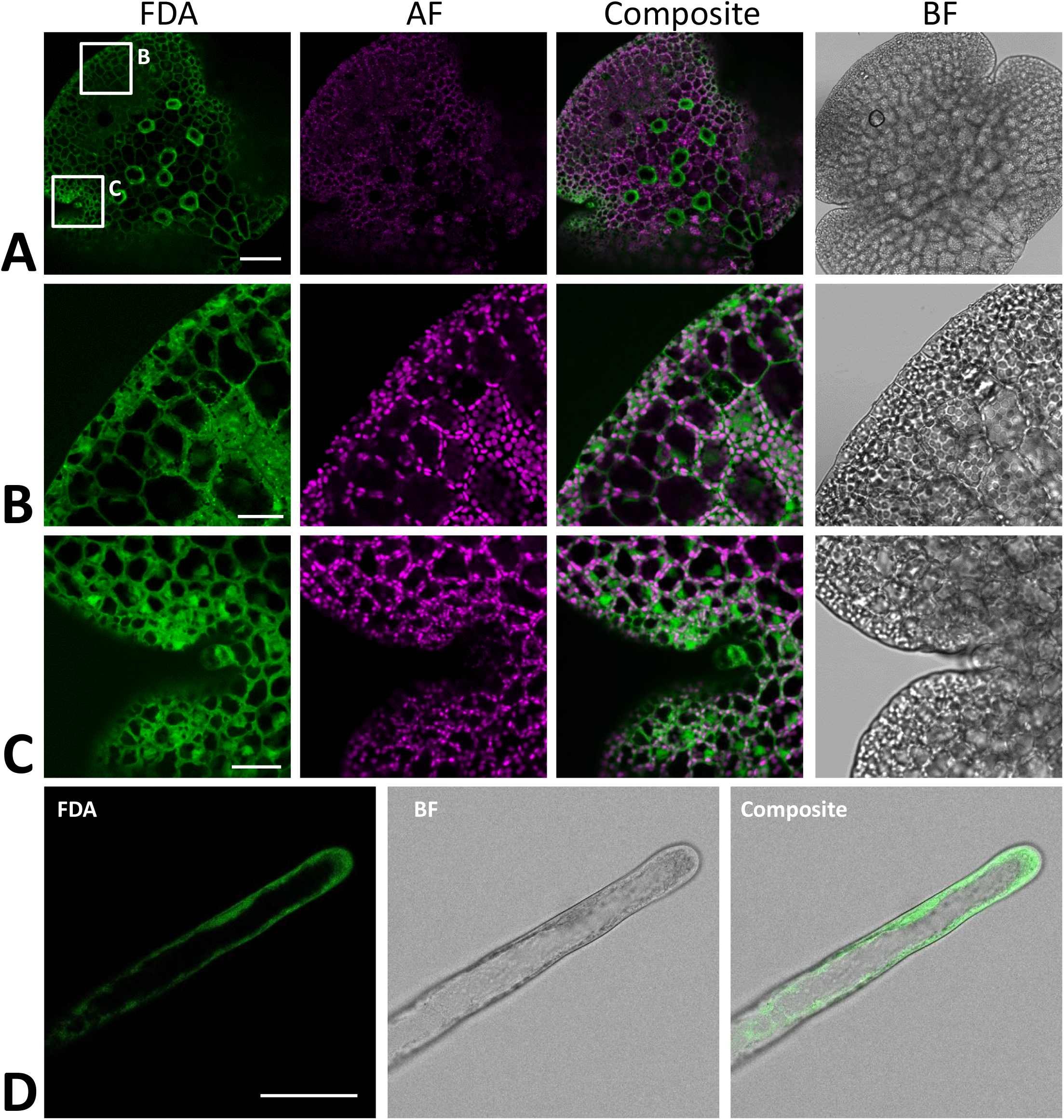
Fluorescein diacetate staining of different *M. polymorpha* cell types. (A) Whole-thallus staining, scale bar = 100 μm, with close-up captures of (B) a distal thallus fragment, scale bar = 30 μm, and (C) a meristematically active apical notch, scale bar = 30 μm. All three images show localization of FDA to the cytoplasm, as contrasted by absence of FDA-specific fluorescence in the vacuole and autofluorescent (AF) chloroplasts. Pictures show maximum projections of z-stack captions (see Materials and methods section for details). (D) FDA staining of a Tak-1 rhizoid of a 5 days-old gemmaling. BF = bright field. Scale bar = 50 μm.

#### Propidium iodide for cell wall staining

Propidium iodide (PI) is an intercalating, red-fluorescent cell dye. It penetrates damaged cell membranes and visualizes nuclei of dead cells by intercalating DNA with low base preference. However, PI cannot pass intact cell membranes and thus is excluded from viable cells, while remaining fluorescent. Therefore, PI can also readily be used to visualize cell wall of living cells. In Arabidopsis, PI is regularly utilized for counterstaining of cell walls (Takano et al., 2002, Ubeda-Tomás, 2008), such as for viability assays, frequently combined with FDA (Shahriari et al., 2010, Kong et al., 2018).

We here show successful PI staining of cell walls of *M. polymorpha* (Fig. 11). Strong fluorescence was observed already after short incubation times of 10 min. PI reliably stained the cell walls of living epidermal cells (Fig. 11A) and rhizoids (Fig. 11B) and thus can reveal cell shape and size. This staining was non-toxic as stained rhizoids kept elongating, thereby revealing the usefulness of PI staining for studying rhizoid tip-growth (Video S1).

**Fig. 11.**
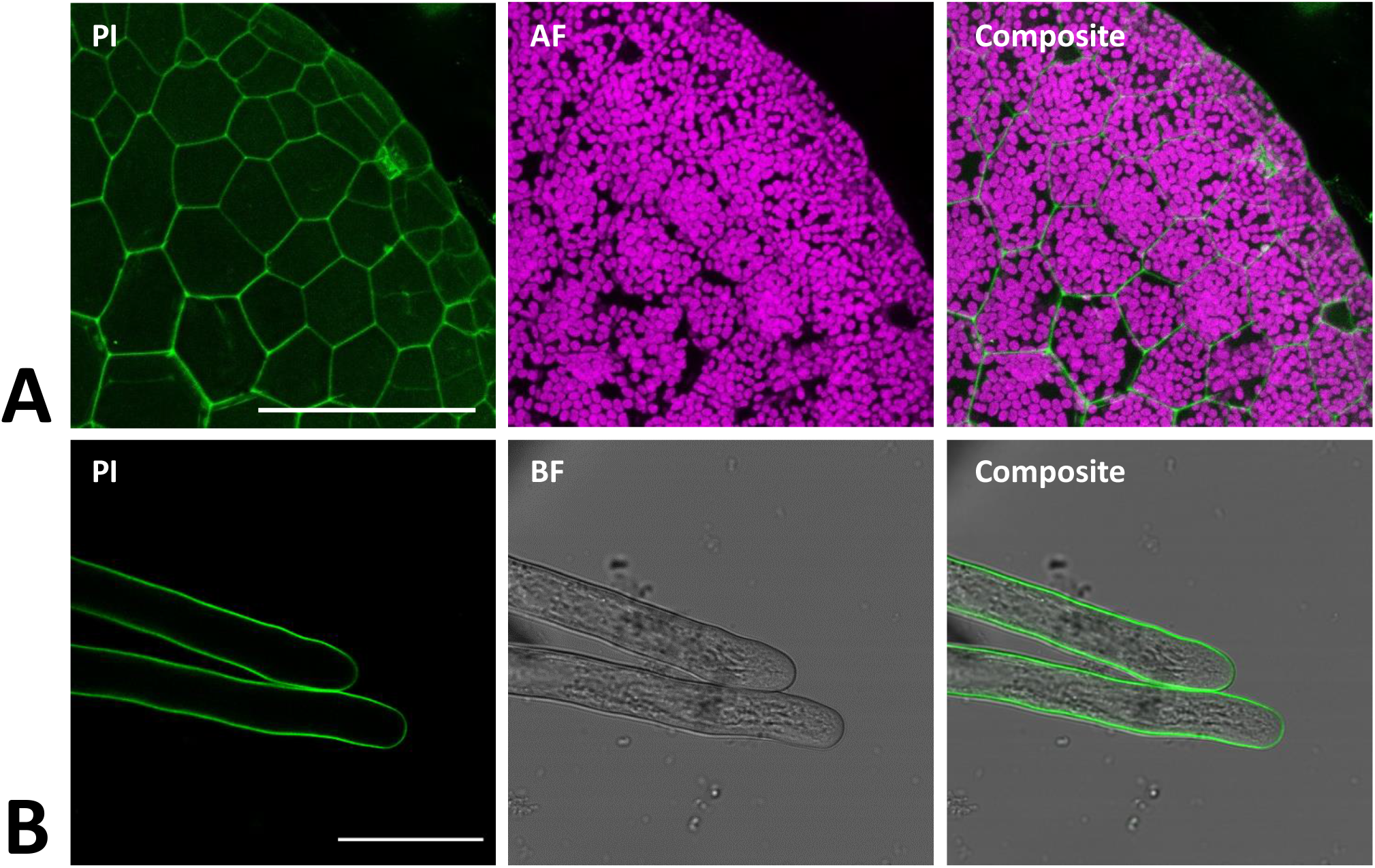
Propidium iodide staining of different *M. polymorpha* cell types. Propidium iodide (PI) staining of Tak-1 rhizoids of a 2 days old gemma, staining the cell wall of both thallus epidermal cells (A; Pictures show maximum projections of z-stack) and rhizoids (B). Scale bars = 100 μm (A) and 50 μm (B). BF = bright field. AF = Autofluorescence (detected at an emission of 680 – 700 nm).

#### Nuclei of M. polymorpha cannot be reliably stained with 4′,6-diamidino-2-phenylindole (DAPI)

DAPI is one of the most common DNA fluorochromes enabling staining and visualization of nuclei of dead but also viable cells, as it is able to pass cell membranes – however often with weak effectiveness. Upon excitation with ultraviolet light, DAPI emits blue fluorescence at a maximum of 461 nm. DAPI binds stoichiometrically to adenine-thymine rich regions of DNA. DAPI also has a weak binding capacity to RNA, however emission is then shifted to 500 nm. Thus, DAPI is frequently utilized not only to visualize nuclei in trichomes, epidermal pavement cells or root cells (Kirik et al., 2001, Spitzer et al., 2006, Lee et al., 2006), but also to quantify DNA content in Arabidopsis, being a reliable tool to discover endoreduplication (Schnittger and Hülskamp, 2007, Bramsiepe et al., 2010, Bhosale et al., 2018). Kondou et al., 2019, report a functional DAPI staining of nuclei in wholemount samples of fixed epidermal cells of *M. polymorpha.* In this study, we tested staining of fixed (i.a. after a modified version of the protocol by Kondou et al., 2019) but also of viable thallus epidermal cells of Marchantia. Surprisingly, despite usage of gemmae at different developmental stages, short to long DAPI incubation periods, preceding and subsequent de-staining steps using EtOH, different methods of fixation (for more details see Material and Methods section), we were unable to stain and visualize nuclei of Marchantia with DAPI (Fig. S3). In our hands, DAPI accumulated on cell walls and to a weaker extent in the cytoplasm but did not enter the nucleus. To demonstrate functionality of the used DAPI solution, we stained Arabidopsis leaves in parallel (Fig. S3), showing strong and distinctive visualization of nuclei. Staining of DNA by PI after fixation also failed in our hands (data not shown). It remains to be elucidated, why nuclei of *M. polymorpha* seem to be hardly accessible to DNA fluorochromes. Until then, we either suggest to use a protein marker localizing in nuclei (*e.g.* AtKRP1) and to generate stably expressing Marchantia lines if needed; or to visualize S-phase nuclei with 5-Ethynyl-2’-deoxyuridine (EdU) staining, as reports show its functionality in *M. polymorpha* (Furuya et al., 2018, Busch et al., 2019)

#### FM4-64 staining for visualization of plasma membrane and endocytic vesicles

The lipophilic steryl dye FM4-64 (3-triethylammoniumpropyl)-4-(6-(4-(diethylamino)-phenyl)-hexatrienyl) pyridinium-dibromide) is commonly used as marker for the outer leaf of the cellular plasma membrane. Staining of young gemmae with FM4-64 resulted in a clear fluorescence signal at the cellular boundaries, likely representing the plasma membrane (Fig. 12A). Upon co-staining with FDA, the FM4-64-specific plasma membrane signal at the cell periphery was clearly distinct from the cytoplasmic FDA signal (Fig. 12B). Altogether, these findings support FM4-64 as a reliable marker dye to label the outer cellular membrane via single or co-staining in Marchantia.

**Fig. 12.**
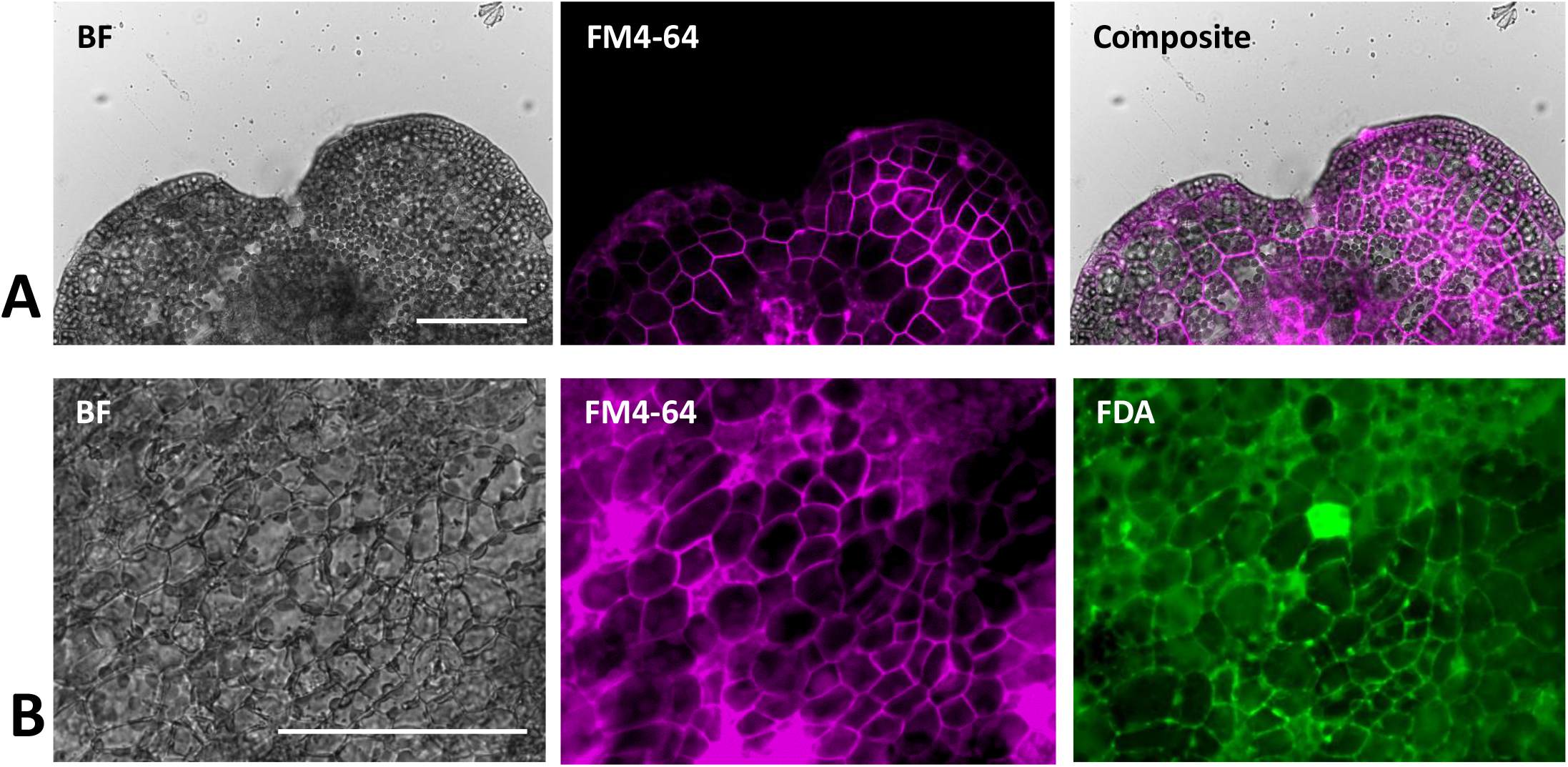
FM4-64 staining of *M. polymorpha* thallus epidermal cells. (A) FM4-64 staining of a 2 days old Tak-1 gemmaling, staining the plasma membrane of thallus epidermal cells. (B) Co-staining of FM4-64 and FDA showing opposing plasma membrane- and cytoplasm-localized fluorescence signal. BF = bright field. FDA = Fluorescein-diacetate. All scale bars = 100 μm.

#### Concluding remarks

We here present a comprehensive and reliable toolkit for visualization of intracellular architecture and dynamics in *M. polymorpha*, an emerging model system used to study land plant evolution. All methods described are based on standard techniques used in other systems and can be executed and analyzed within 1 −2 working days, therefore allowing time-efficient analysis of basic intracellular traits, such as organelle organization and cell architecture, both in fixed and viable cells. The possibility to mark viable cells additionally allows their analysis in live-imaging setups, as we demonstrate with growing rhizoids stained with PI. A comprehensive list of transiently expressed markers covering the majority of intracellular organelles and structures, allows fast assessment of aforementioned intracellular dynamics in viable cells, but also provides a quick possibility for initial tests of functionality and correct localization of cloned fluorescent constructs before committing to comparatively time-costly stable plant transformation. Finally, we demonstrate the BiFC system to be functional in Marchantia epidermal cells, thus representing a quick and straightforward technique to test for protein-protein interactions *in vivo*, which should be confirmed with other protein-protein interaction assays such as Yeast-2-Hybrid-like, FRET-FLIM and protein pulldown approaches. Altogether, we provide a series of quick and useful techniques to exploit the potential of an emerging model system to the maximum extent possible.

## Acknowledgements

We thank Dr. Marc Jakoby for providing aliquots of the AtKRP, SKL motif, AtNPSN12, AtSYP32 and AtGot1p homolog vectors. We thank Dr. Alexandra Steffens for providing aliquots of the AtDCP1 and AtDCP2 expression vectors. We thank Dr. Lisa Stephan for providing aliquots of the AtMYC1 and AtTTG1 BiFC vectors. We thank Dr. Clement Champion and the research group of Prof. Liam Dolan (University of Oxford) for provision of an aliquot of the PM marker vector MpSYP13a. We thank Dr. Joachim F. Uhrig for donation of pCL112/113 vectors. This research was partly funded by a short-term stipend of the Deutscher Akademischer Austauschdienst (DAAD) to J.W.; the University of Cologne, and grant from the University of Cologne Centre of Excellence in Plant Sciences to A.B.-D.

## Author contributions

J.W., E.K., M.H. and A.B.-D. conceived the experiments.

J.W., E.K., R.L. and A.B.-D. performed the experiments.

J.W., E.K. and A.B.-D analyzed the data.

J.W. and E.K. wrote the manuscript with contributions of M.H. and A.B.-D.

---

**Tab. S1:**
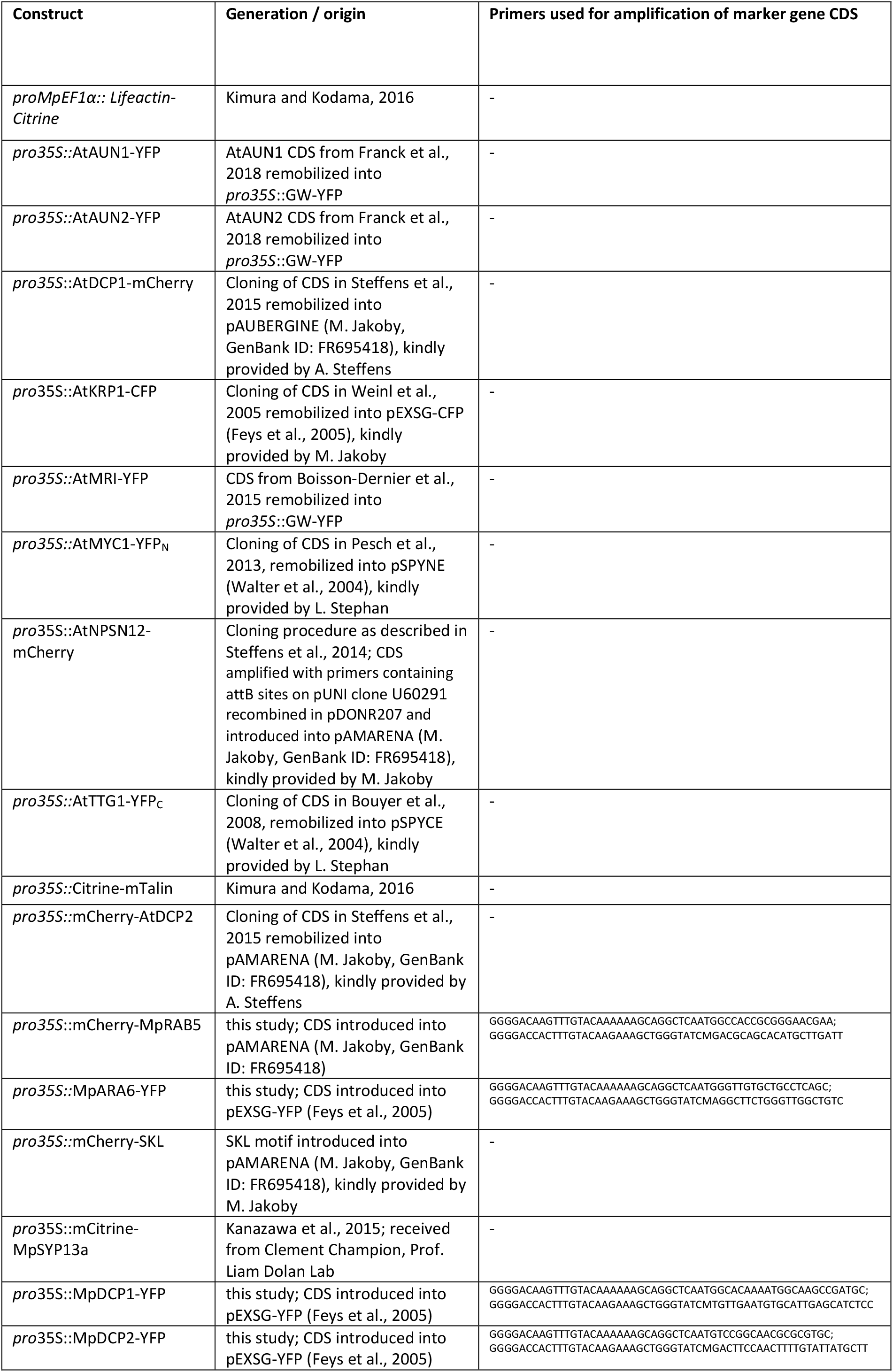
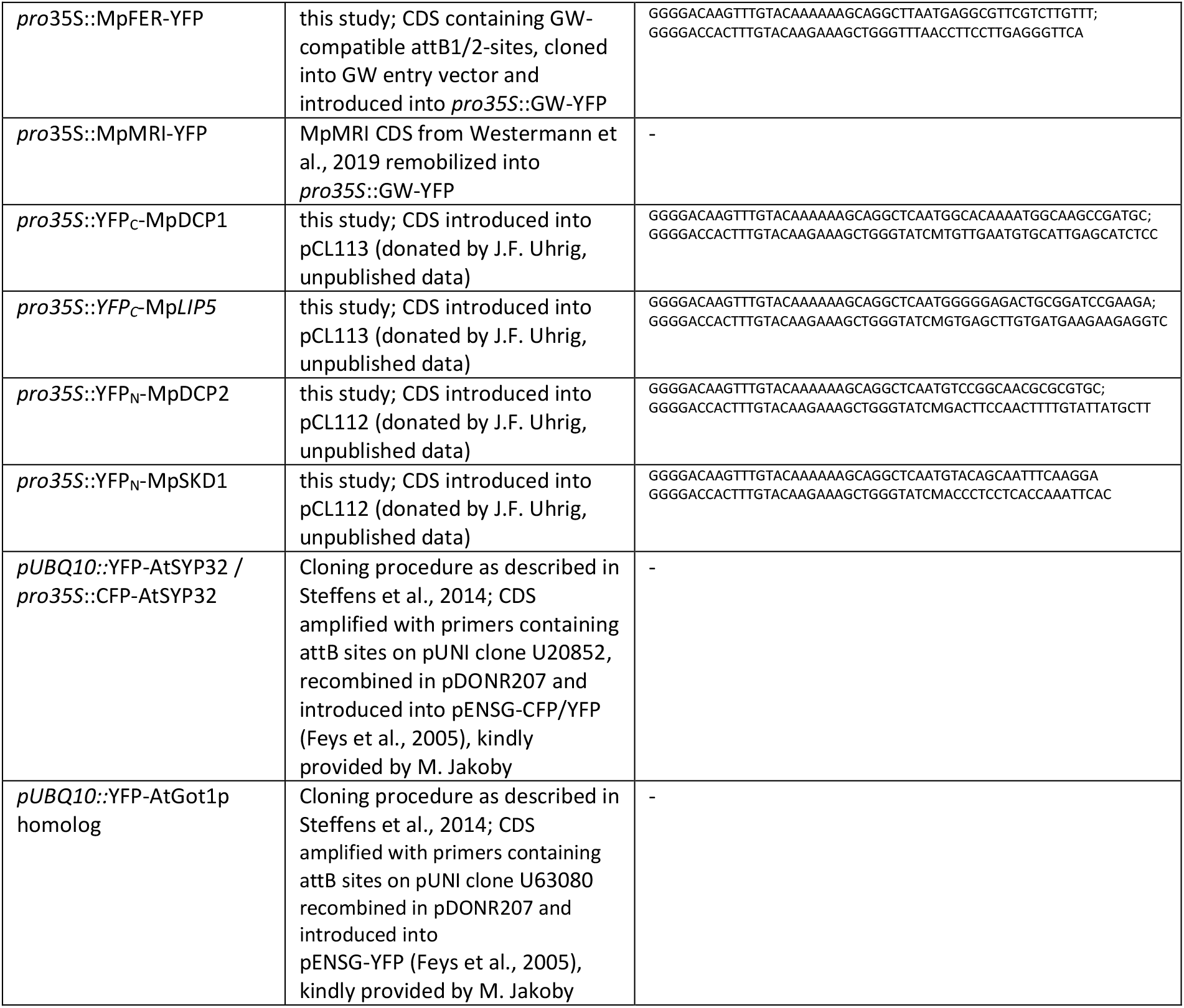
Comprehensive list of all marker constructs used for biolistic transformation. The list includes their origin (referenced publication or own generation), as well as the oligonucleotide sequences used as primers for amplification of new marker gene CDS. GW = Gateway-compatible cassette.

**Tab. S2:** Excitation and captured emission wavelengths used for analysis of fluorescent markers.

| Fluorophore | Excitation | Captured Emission |
| --- | --- | --- |
| CFP | 458 nm | 470 nm – 480 nm |
| YFP | 514 nm | 524 nm – 530 nm |
| mCherry | 561 nm | 607 nm – 618 nm |
| FDA | 514 nm | 500 nm – 540 nm |
| PI | 561 nm | 610 nm – 630 nm |
| DAPI | 405 nm | 450 nm – 470 nm |

**Tab. S3:** Quantification of co-bombardment efficiency in *M. polymorpha* biolistic transformation. Data from 9 independent co-transformation events of marker proteins used in this study. Scans of at least two transformed cells were used for the quantification.

| Cells expressing both marker proteins | Total number of transformed cells | Efficiency of co-transformation [%] |
| --- | --- | --- |
| 2 | 2 | 100 |
| 8 | 12 | 67 |
| 2 | 2 | 100 |
| 1 | 2 | 50 |
| 2 | 2 | 100 |
| 2 | 2 | 100 |
| 3 | 3 | 100 |
| 1 | 2 | 50 |
| 1 | 2 | 50 |
| 5 | 8 | 63 |
| 4 | 5 | 80 |
| 1 | 2 | 50 |
| 1 | 2 | 50 |
| 3 | 3 | 100 |
| 4 | 6 | 67 |
| 7 | 7 | 100 |
| 1 | 2 | 50 |
| 2 | 2 | 100 |
| 1 | 2 | 50 |
| 1 | 2 | 50 |
| 3 | 4 | 75 |
| 2 | 2 | 100 |
| 3 | 5 | 60 |
| **SUM: 60** | **SUM: 81** | **Average: 74** |
|  |  | SD: 23 |

**Fig. S1:**
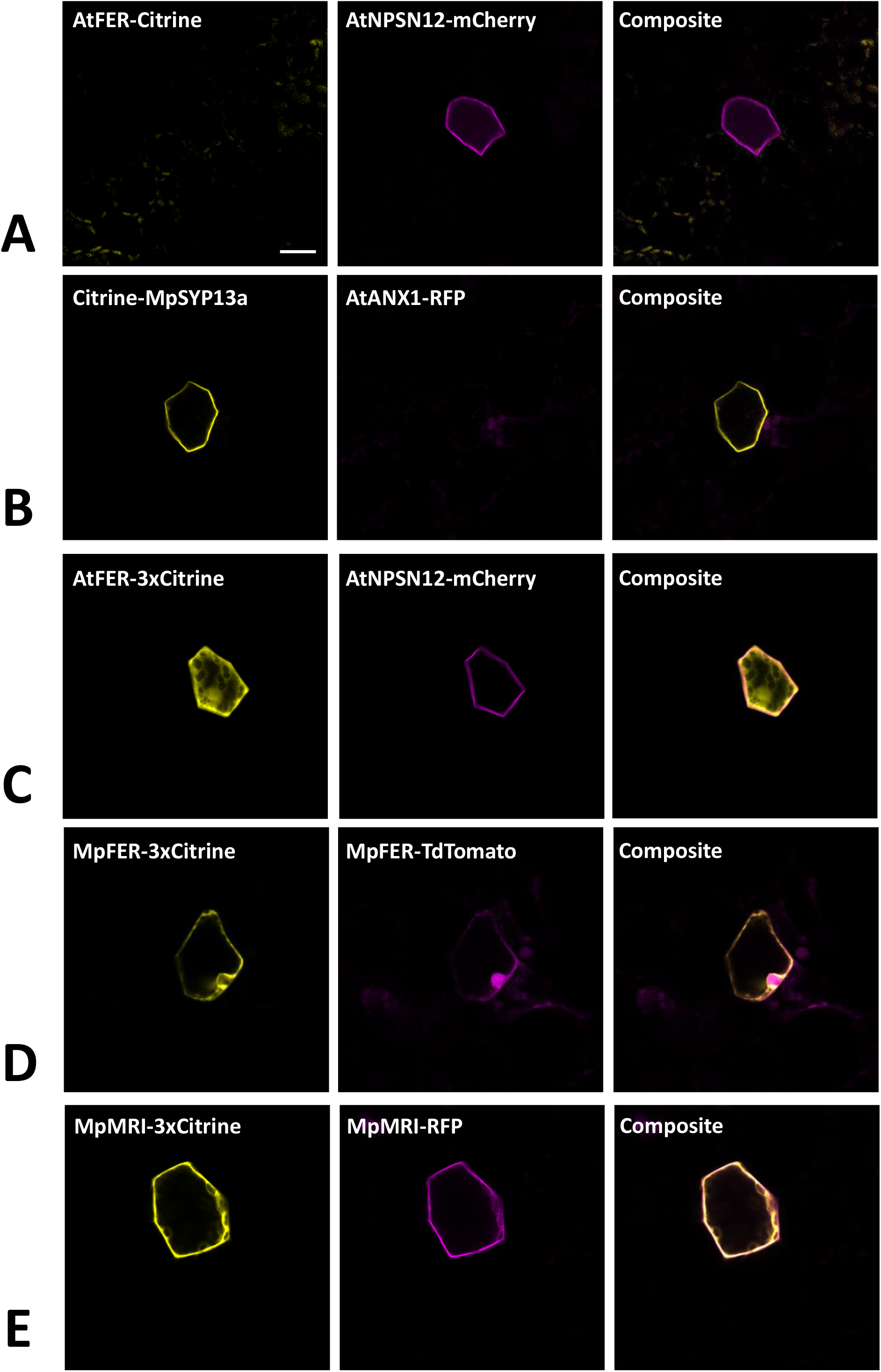
Co-expression of MLRs and MRI with single or triple tags in Marchantia epidermal cells. (A) and (B) Arabidopsis MLRs fused to single fluorescent tag are not expressed. (C) and (D) the 3xCitrine tag leads MLRs to localize to the cytoplasm. (E) the 3xCitrine tag leads to normal cytosolic and plasma membrane localization of MpMRI. Pictures show maximum projections of z-stack captions (see Materials and methods section for details). Scale bar = 20 μm.

**Fig. S2:**
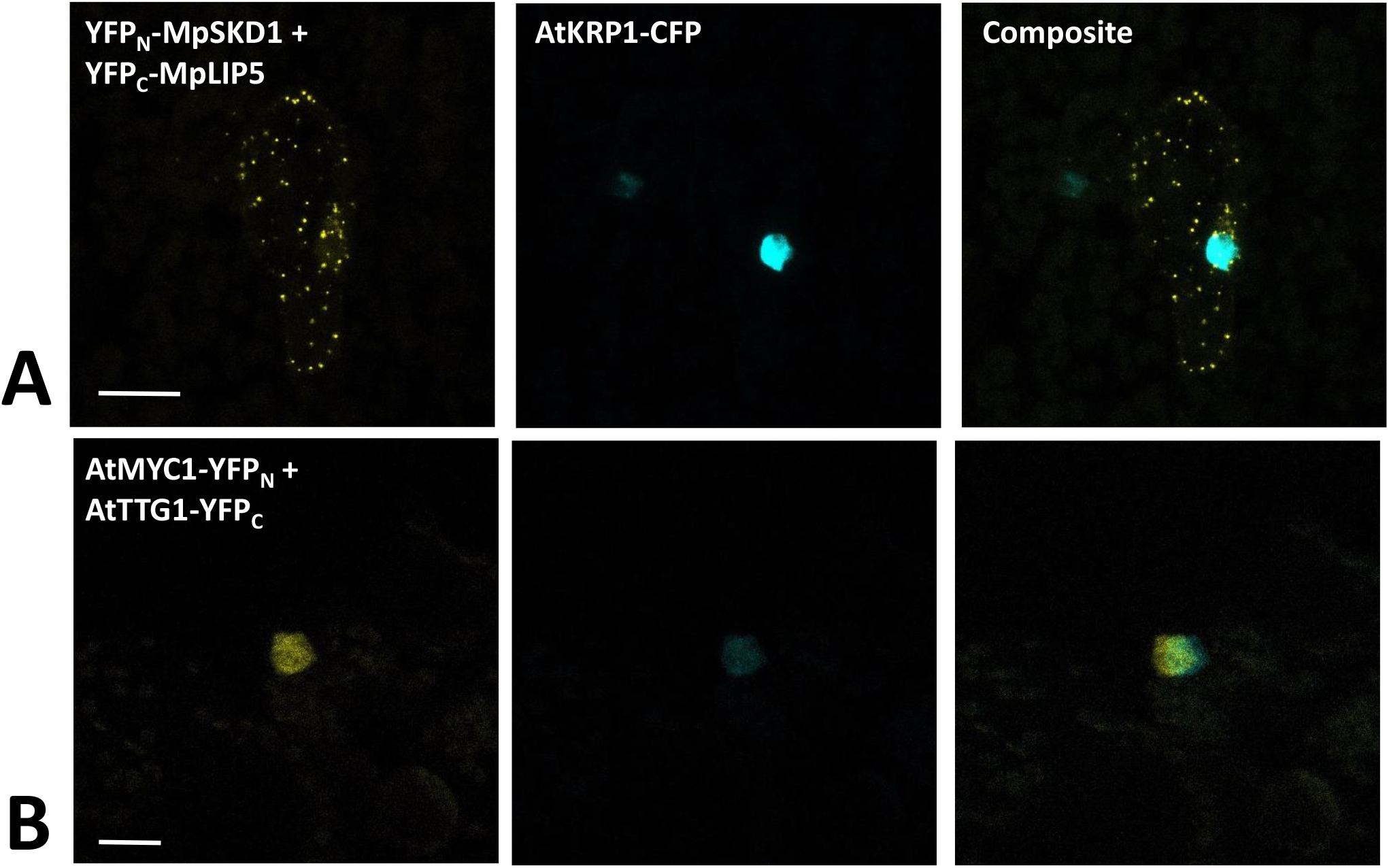
Bimolecular fluorescent complementation assay quality controls. (A): The functionality of the negative control MpLIP5 was confirmed via co-bombardment of split-versions of MpLIP5 and the Marchantia homolog of the known Arabidopsis LIP5 interactor MpSKD1, showing a clear protein interaction in dot-like foci. (B): Split-YFP fusion constructs of AtMYC1 and AtTTG1, known interactors, were co-bombarded and shown to physically interact in *M. polymorpha* thallus epidermal cells, supporting the functionality of AtMYC1-YFP_N_. The constructs were co-bombarded with nuclear marker AtKRP1. Scale bar = 20 μm. Pictures show maximum projections of z-stack captions (see Materials and methods section for details).

**Fig. S3:**
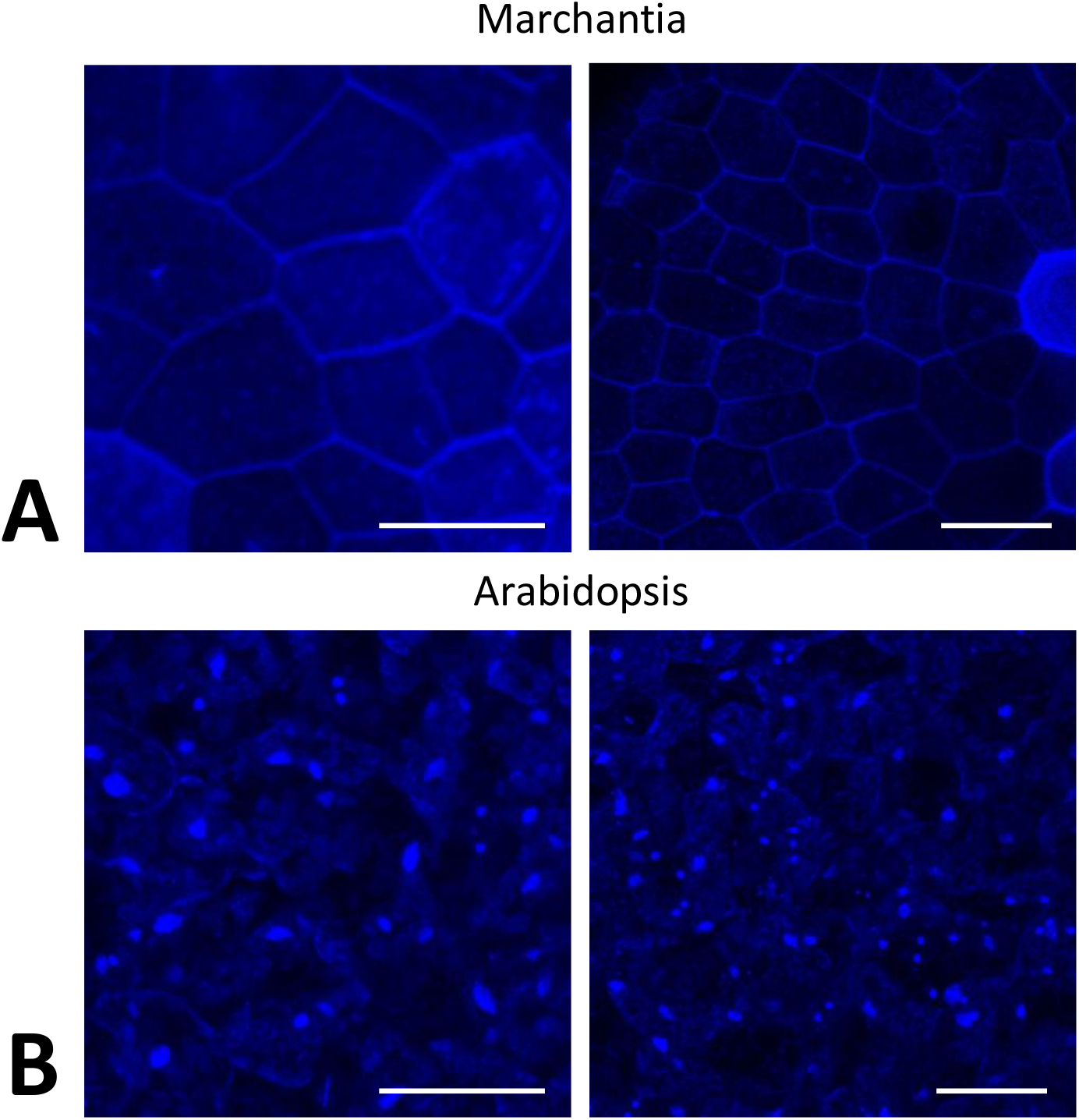
Nuclei of *M. polymorpha* cannot be readily stained with DAPI. (A) DAPI staining of Tak-1 epidermal cells of a 4 days-old gemmaling. (B) DAPI staining of leaf epidermal cells of a 2 weeks old *A. thaliana* plant. Note the stained nuclei. All scale bars = 50 μm.

**Video S1: Growing rhizoids stained with propidium iodide.**

